# Uncovering the temporal dynamics and regulatory networks of thermal stress response in a hyperthermophile using transcriptomics and proteomics

**DOI:** 10.1101/2023.05.02.539125

**Authors:** Felix Grünberger, Georg Schmid, Zubeir El Ahmad, Martin Fenk, Katharina Vogl, Robert Reichelt, Winfried Hausner, Henning Urlaub, Christof Lenz, Dina Grohmann

## Abstract

Extremophiles, such as the hyperthermophilic archaeon *Pyrococcus furiosus*, thrive under extreme conditions and must rapidly adapt to changes in the physical parameters of their natural environment for short-term and long-term survival. When inhabiting hydrothermal vents, these organisms face substantial temperature gradients, necessitating the evolution of adaptive thermal stress mechanisms. However, the dynamics and coordination of cellular responses at the transcriptome and proteome levels remain underexplored. This study presents an integrated analysis of RNA-sequencing and mass spectrometry data to elucidate the transcriptomic and proteomic responses to heat and cold shock stress and recovery in *P. furiosus*. Our results reveal surprisingly rapid and dynamic changes in gene and protein expression patterns associated with these stress responses. Heat shock triggers extensive transcriptome reprogramming, orchestrated by the transcriptional regulator Phr, which targets a broader gene repertoire than previously demonstrated. For heat shock signature genes, RNA levels swiftly return to baseline upon recovery, while protein levels remain persistently upregulated, reflecting a rapid but more sustained response. Intriguingly, cold shock at 4°C elicits distinct short-term and long-term responses at both RNA and protein levels. By conducting a cluster analysis, we identified gene sets with either congruent or contrasting trends in RNA and protein changes. Notably, these clusters represent well-separated arCOG groups and appear to be tailored to their individual cellular responses. Our study provides a comprehensive overview of the cellular response to temperature stress, advancing our understanding of stress response mechanisms in hyperthermophilic archaea and provide valuable insights into the molecular adaptations that facilitate life in extreme environments.

## Introduction

The ability of extremophilic microorganisms to adapt and thrive in extreme environments has captivated the scientific community for decades (1–3). These organisms provide unique opportunities to investigate the molecular basis of stress response, adaptation and recovery of cellular activities, offering valuable insights into fundamental biological processes and enabling novel biotechnological applications (4). Among extremophiles, the hyperthermophilic archaeon *Pyrococcus furiosus* has become a model organism for studying the molecular strategies employed by thermophiles to withstand high temperatures (5–7). It is a marine-living anaerobic organism that grows over a broad temperature range from 70 to 104°C and can be found in black smokers within hydrothermal vent systems that are characterized by significant temperature gradients ranging from 2°C to 400°C (Fig. S1 A) (8–11). Since hydrothermal vents are sterile during their initial formation, it was first hypothesized and later experimentally shown that they are colonized by hyperthermophilic archaea from the surrounding 4°C-cold seawater (12, 13). Moreover, evidence suggests that *Pyrococcus* must possess mechanisms to withstand extended cold-shock periods as it has been successfully recultivated from cooled-down submarine plumes and floating volcanic slick taken over one kilometre away from the active zone of an erupted submarine volcano (14). This observation has also been replicated in the laboratory, where it was demonstrated that hyperthermophiles could survive for at least nine months in cold environments and react within seconds upon returning to their optimal growth temperature by initiating motility (15).

In general, hyperthermophilic archaea have adapted to thrive at temperatures exceeding 80°C through various molecular mechanisms, including stabilization of tRNAs and rRNA by higher GC-content, enrichment in hydrophobic and charged amino acids, alterations in protein structure, unique membrane composition, increased investment in nucleoid-associated proteins and positive DNA supercoiling by reverse gyrase (16–20). However, the biomolecular and biophysical challenges associated with heat shock above the individual temperature limit of each organism are at least to some extent shared across the domains of life (21, 22). In contrast, less is known about shared molecular principles dealing with cold shock response, especially in archaea (23, 24).

Extreme temperature fluctuations pose significant challenges to cellular macromolecules, such as proteins, nucleic acids, and lipids. For instance, elevated temperatures can lead to protein denaturation, aggregation, and loss of function during heat shock. Concurrently, DNA and RNA can undergo structural alterations, impairing replication, transcription, and translation processes (25). On the other hand, cold shock induces the stabilization of secondary structures in nucleic acids, impeding their proper function and reducing transcription and translation efficiency. Furthermore, low temperatures affect membrane fluidity, potentially impairing membrane-associated processes and transport (23, 24, 26).

In bacteria, the heat shock response is primarily regulated by the conserved sigma factor σ32, which promotes transcription of heat shock genes encoding chaperones, such as DnaK and GroEL, and proteases responsible for protein refolding and degradation (27). In addition to the positive regulation by sigma factors, it is well-known that transcriptional repressors, like HrcA or HspR, can also induce expression of heat shock genes by dissociating from the promoter (28–30). Cold shock responses involve the synthesis of cold-induced proteins, such as CspA in *Escherichia coli*, which counteract the effects of low temperatures on nucleic acids and cellular processes. These Csps are small, nucleic acid-binding proteins that are widely distributed in bacteria and structurally highly conserved, containing a cold shock domain that enables binding to target RNA and DNA. Also, they function as RNA chaperones, destabilizing secondary structures at low temperatures and facilitating transcription and translation (24, 31, 32).

In eukaryotes, the heat shock response is orchestrated by heat shock transcription factors, which modulate the expression of heat shock proteins like Hsp70 and Hsp90. On the other hand, the cold shock response involves diverse mechanisms, such as changes in membrane lipid composition and synthesis of cold-inducible RNA-binding proteins (33–36).

Organisms across the tree of life are challenged by stressors such as heat and cold shock. The specific molecular players and regulatory networks involved in these processes, however, can differ significantly. For example, cold shock domain proteins are absent in all thermophilic and hyperthermophilic archaea and the regulation of the cold shocks response in archaea is less well understood compared to bacteria and eukaryotes (37). However, for the heat shock response some regulatory mechanisms have been identified, such as the transcription factor (TF) Phr in *P. furiosus*, which recognizes a palindromic DNA sequence and acts as a negative regulator of many heat-inducible genes (38–40). In contrast, no thermal-responsive regulator has been identified in Crenarchaeota and thus is also not present in the well-described thermophile *Sulfolobus acidocaldarius* (21, 41). Nevertheless, *S. acidocaldarius* exhibits a classic heat shock response characterised by the induction of small chaperones and the thermosome. Notably, its genome encodes two to three different thermosome subunits, and their assembly is modulated based on the prevailing environmental conditions (21, 42).

In addition to transcription factor-based regulation, archaeal-specific small RNAs (asRNAs) and RNA-binding proteins like proteins of the Sm-like protein family have been implicated in fine-tuning regulation of stress responses on the post-transcriptional level (43–46).

Despite significant advancements in the field, our understanding of the complex regulatory networks and adaptive mechanisms employed by *P. furiosus* in response to extreme temperature variations remains limited, especially when considering the time-related aspects of these processes on the RNA and protein level.

This study aims to unravel the temporal dynamics of thermal stress response in the hyperthermophilic archaeon *P. furiosus* using an integrative omics approach that combines transcriptomic and proteomic analyses at multiple time points. Our objective is to analyse shifts in gene expression and protein abundance that occur as the organism experiences conditions mimicking the thermal environment of hydrothermal vents. To achieve this, we first compare our findings with the established knowledge of responses on the transcriptional level and mechanisms of the heat and cold shock response. We furthermore integrate the transcriptomic and proteomic data to generate a more comprehensive understanding. Moreover, we consider the temporal resolution of our experimental setup to identify gene clusters and key cellular processes that display similar regulatory patterns. Furthermore, we explore potential regulatory elements, such as promoters, terminators, and operons, that may influence the transcriptomic landscape under stress conditions. In particular, we seek to enhance our understanding of the temporal resolution of Phr-regulated heat shock response. Ultimately, this in-depth analysis aims to expand our knowledge of the molecular adaptations that enable long-term survival in cold seawater and recolonization of black smokers, contributing to the broader understanding of extremophile biology.

## Results

### Experimental setup and analysis strategy for studying the heat and cold shock response in Pyrococcus furiosus

To investigate the temporal dynamics of transcriptomic and proteomic responses of *P. furiosus* to heat and cold shock, we designed a custom temperature-controlled system for rapid cooling and reheating of samples (Fig. S1 B). Cells were initially cultivated at an optimal temperature of 95°C to mid-exponential growth phase, serving as the control condition (Ctrl). Cold shock (CS) was simulated by shifting the temperature to 4°C, with samples taken at 20 min (CS 1), 2 h (CS 2), and 24 h after transfer of the samples to 4°C (CS 3) (Figure 1 A). Heat shock (HS) samples were collected after 5 min (HS 1) and 15 min (HS 2) after shifting the temperature to 105°C, a temperature that inhibits cell growth (8). In both setups, recovery samples (CS R, HS R) were obtained by returning cells to their optimal growth temperature following the final shock time point.

**Figure 1.**
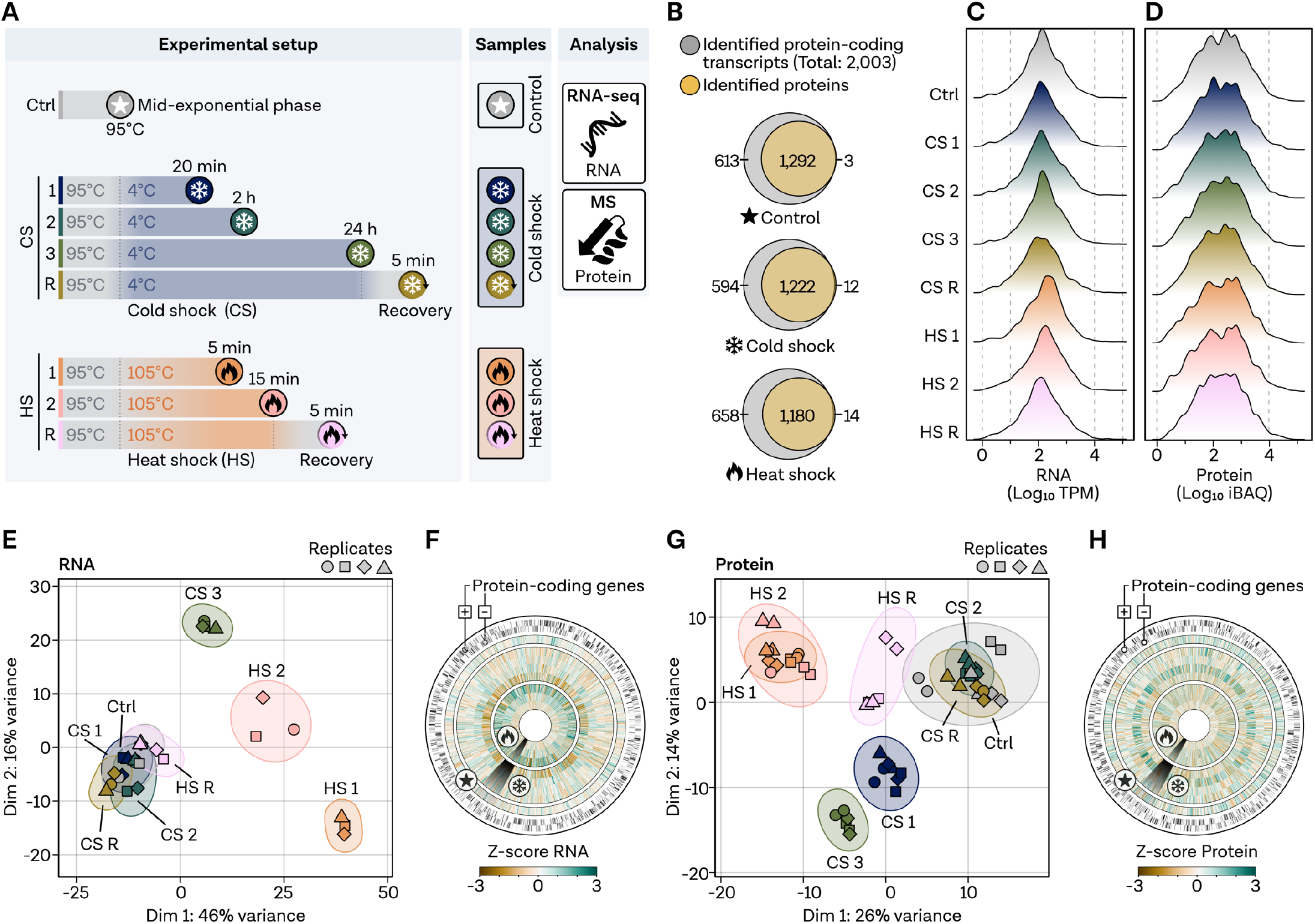
Validation of the experimental setup by temporal analysis of the heat and cold shock response in *Pyrococcus furiosus* via transcriptomics and proteomics. **A,** Experimental design illustrating the seven stress-related conditions, including cold shock (CS) at 4°C and heat shock (HS) at 105°C, along with recovery samples, analyzed using RNA-seq and proteomics, compared to the control condition (Ctrl) at the optimal growth temperature of 95°C. **B**, Venn diagram displaying the overlap between identified protein-coding transcripts (grey) and total identified proteins (yellow) under stress-related and control conditions. Genes only identified using RNA-seq or proteomics are shown at the left and right side, respectively. **C**, Distribution of transcripts per million (TPM) normalized transcript counts, color-coded based on the legend in panel A. **D**, Distribution of intensity based absolute quantification (iBAQ) values for proteomics samples. **E**, Principal component analysis (PCA) of RNA-sequencing samples based on total counts; replicates are indicated using different symbols, with outliers removed. **F**, Circular genome-wide plot of the *P. furiosus* genome (position 0 at top) with protein-coding genes on the two outer rings; Z-score normalized TPM counts color-coded by dark brown (negative) to dark green (positive) for each condition in the following order (outer to inner): Ctrl, CS 1, CS 2, CS 3, CS R, HS 1, HS 2, HS R. **G**, PCA and **H**, genome wide circular plot for mass spectrometry samples analogous to panels E,F.

We performed deep RNA sequencing (RNA-seq) on rRNA-depleted libraries, yielding approximately 600-fold transcriptome coverage and detecting 95% (Ctrl), 91% (CS) and 92% (HS) of all 2,003 protein-coding transcripts in *P. furiosus* (Figure 1 B) (counts listed in Table S1) (47). Additionally, we conducted mass spectrometry (MS) analysis on all samples, detecting 65% (Ctrl), 62% (CS) and 60% (HS) of the proteins, respectively. iBAQ values were used as accurate proxies for protein abundance. Nearly all proteins detected by MS with an IBAQ value of ≥10 had corresponding transcripts with at least 10 transcripts per million (TPM) RNA counts (Figure 1 B,C,D). Uniform distribution of TPM-normalized RNA-seq counts and iBAQ values suggests high data quality, allowing for comparisons between conditions and methods (Figure 1 C,D).

Principal component analysis (PCA) revealed distinct temporal and temperature-dependent responses at the RNA level, with the most significant variance observed after HS 1, HS 2, and CS 3 (Figure 1 E). To determine whether differences in the PCA can be explained by stress islands in the genome, we visualized z-score normalized TPM counts globally, finding that transcript changes were dispersed throughout the transcriptome (Figure 1 F). Variance at the RNA level was corroborated at the protein level, with two exceptions (Figure 1 G): First, the HS recovery sample did not cluster with the Ctrl condition but was positioned between HS 1 and the control, suggesting a prolonged protein response. Second, CS 1 formed a separate group at the protein level, reflecting substantial variation in protein expression patterns. Similar to the RNA level, proteome changes during CS and HS were not restricted to specific islands (Figure 1 H).

### Transcriptome analysis reveals extensive reprogramming during HS and moderate overlapping response with the CS response in P. furiosus

Having verified the robustness of our experimental design for analyzing temporal aspects of thermal stress and recovery responses in *P. furiosus*, we proceeded with our investigation in two stages. Therefore, we first conducted a comprehensive transcriptome analysis to validate the selected time points and temperatures based on known stress responses and to identify potential regulatory aspects before integrating the results with the proteomics data. A 5-minute HS at 105°C (HS 1) induced significant changes in 68% of the transcriptome, with 330 genes upregulated over 2-fold and 411 genes downregulated over 2-fold (Figure 2 A). Notably, several upregulated genes included well-known HS proteins, confirming the effectiveness of our experimental setup. Specifically, HS in *P. furiosus* has been shown to induce the expression of the thermosome (log_2_FC: 2.7), small chaperones, such as HSP20 (log_2_FC: 5.4), VAT1 (log_2_FC: 6.7) and VAT2 (log_2_FC: 0.2), and proteases that bind, refold or degrade misfolded proteins accumulating in cells (7) (Figure 2 B). Moreover, we could confirm previous data showing that the synthase (Myo-inositol-1-phosphate synthase: Myo-Synthase, PF1616 4.0) catalyzing a precursor of the compatible solute Di-myo-inositol-phosphate (DIP) accumulates during HS. DIP is suggested to have a protein-stabilizing role in hyperthermophiles, while proteins typically involved in controlling the thermal damage of the proteome, like the ATP-independent chaperone prefoldin (log_2_FC: −1.1), are not upregulated during HS (48, 49). Some of the aforementioned proteins are under the control of the negative transcriptional regulator Phr. We confirmed that all currently known target genes, except PF1292, are substantially upregulated under our selected conditions (40). While the expression of the only ATP-dependent protease in *P. furiosus*, the proteasome (alpha-subunit log_2_FC: −0.4; beta-subunit log_2_FC: 0.2), is not increased during HS, the proteasome-assembling chaperone homolog PbaA (log_2_FC: 3.9) that forms a complex with PF0014 (log_2_FC: 3.9) is highly upregulated (7, 50) (Figure 2 B).

**Figure 2.**
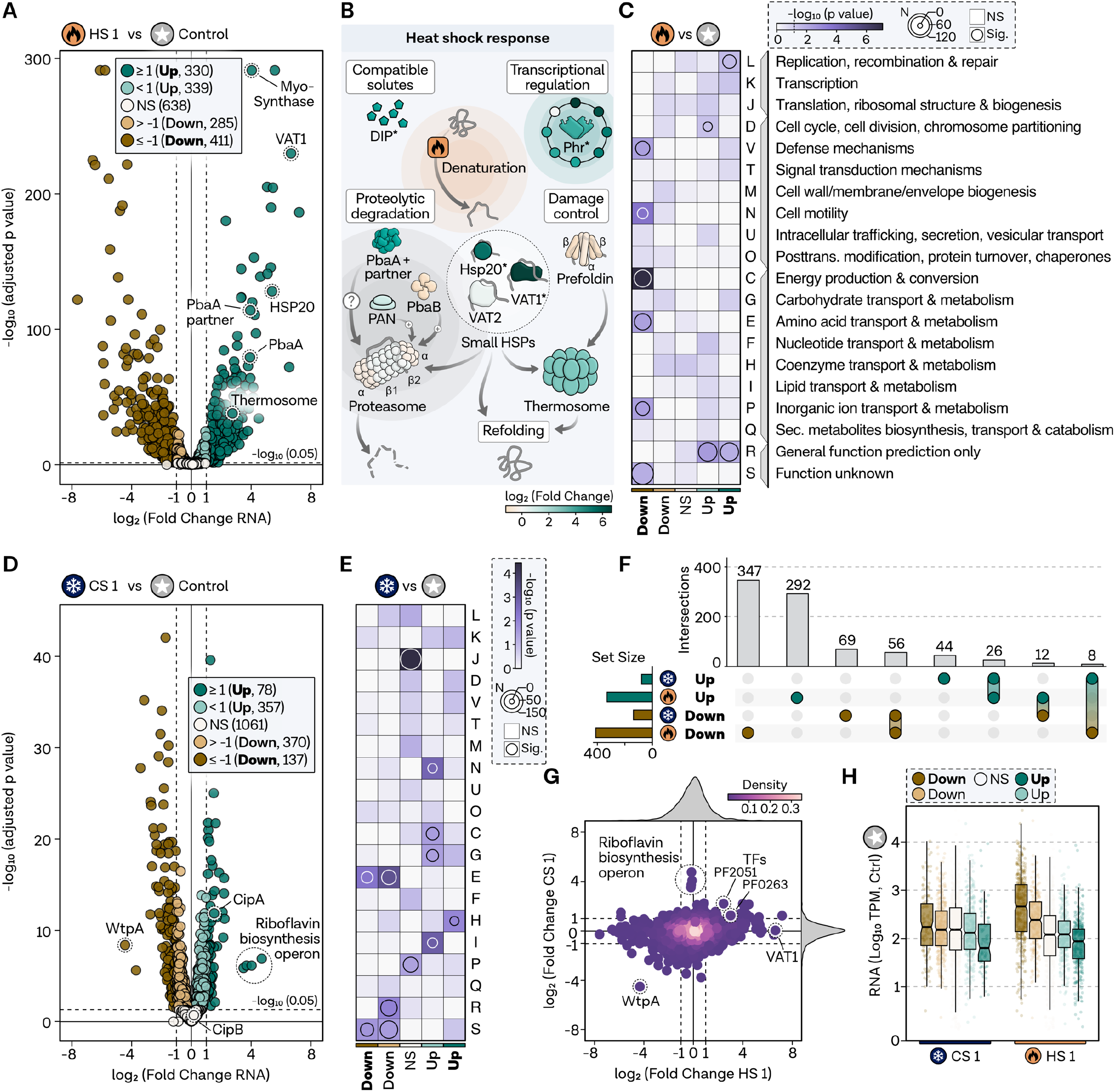
HS leads to extensive reprogramming on the transcriptome level with moderate overlap to CS response. **A**, Volcano plot displaying log_2_ fold changes (x-axis) and significance (−log_10_ adjusted p-value, y-axis) comparing transcriptome changes at HS 1 (5 minutes) to the control condition. Protein-coding transcripts are categorized by significance and fold changes, color-coded as strongly upregulated (padj < 0.05 and log_2_FC >= 1, dark green, bold font), upregulated (padj < 0.05 and log_2_FC < 1 and log_2_FC > 0, light green, normal font), non-regulated genes (NS, padj >= 0.05, white), strongly downregulated (padj < 0.05 and log_2_FC <= −1, dark brown, bold font), downregulated (padj < 0.05 and log_2_FC > −1 and log_2_FC < 0, light brown, normal font). HS genes are highlighted. Significance level of 0.05 is shown as a dashed line. **B**, Schematic representation of the HS response in *P. furiosus*, with genes color-coded according to transcriptome fold changes. **C**, Gene set enrichment analysis of archaeal clusters of orthologous genes (arCOGs), with significance levels indicated by a color bar ranging from white to dark pink. Genes with a p-value < 0.05 are considered significantly overrepresented and highlighted by a circle, with circle size reflecting the number of genes in the category. **D**, Volcano plot showing log_2_ fold changes for CS 1 condition compared to the control condition, with the y-axis displaying significance. A cutoff value of adjusted p-value of 0.05 is indicated by a dashed line, and relevant genes are highlighted. **E**, Functional enrichment analysis using arCOG descriptions; significance levels and the number of regulated genes are shown in the upper legend. **F**, Overlap between strongly upregulated (dark green) and downregulated (dark brown) genes in HS 1 and CS 1 conditions; total gene numbers are shown in horizontal bar graphs, and comparison set numbers are displayed in vertical bar graphs. **G**, Scatter plot comparing the log_2_ fold changes for HS 1 (x-axis) and CS 1 (y-axis), color-coded according to plotting density; fold change densities for each condition are displayed in the side density plots. **H**, TPM normalized expression values from the control condition for respective regulatory groups from panels A and D for CS 1 and HS 1 conditions; box edges delineate the 1st and 3rd quartiles, center line represents the median, and whiskers denote points within 1.5x of the interquartile range.

For systematic HS response analysis, we conducted functional enrichment analysis based on archaeal clusters of orthologous groups (arCOGs) (Figure 2 C) (51). Genes related to replication, recombination, and repair (category L) are overrepresented among highly upregulated genes. In contrast, genes associated with energy production (category C), motility (category N), cell defence (category V), inorganic ion transport (category P), amino acid transport (category E) and unknown groups are overrepresented among highly downregulated genes. The remaining significantly regulated genes with minor fold changes smaller than two do not exhibit clear categories, except for cell cycle overrepresentation in the upregulated group.

In comparison to HS, much less is known about the CS reaction in *Pyrococcus*. Our experimental setup significantly differs from previous analyses, which employed a temperature at which *Pyrococcus* can still grow (70°C), thus limiting direct comparisons (52). Upon immediate CS, 47% of the transcriptome underwent significant alterations (Figure 2 D). However, fewer protein-coding transcripts were strongly affected during CS 1 compared to HS 1, with only 78 strongly upregulated and 137 strongly downregulated genes observed. The most substantial upregulation occurred for a riboflavin-synthesis operon (PF0061-PF0064, log_2_FC: 4.5), potentially under the control of the transcriptional regulator RbkR (PF0988) (53). Among the cold-responsive solute-binding proteins CipA (log_2_FC: 1.5) and CipB (log_2_FC: 0.2), only the former was induced significantly during CS (52). Functional enrichment analysis revealed that genes related to coenzyme transport and metabolism were overrepresented in highly upregulated genes during CS (Figure 2 E). Additionally, categories C and G, corresponding to energy production and carbohydrate transport, and lipid transport (category I) were overrepresented in upregulated genes. In contrast, genes associated with amino acid transport and metabolism were downregulated on the RNA level during CS. Interestingly, genes involved in translation and ribosome function, as well as inorganic ion transport, remained unaffected at the RNA level. The functional overlap between groups of regulated genes of the HS and CS response was limited, except for the shared downregulation of amino acid metabolism.

To further investigate this, we compared the gene-specific regulation of groups that are highly up- or downregulated under both conditions (Figure 2 F). Almost half of the genes downregulated during CS were also downregulated under HS, and this pattern was consistent for upregulated genes as well (Fig. S2). Two currently uncharacterized transcription factors were upregulated under both CS and HS, suggesting a potential role in stress regulation (Figure 2 G). While the riboflavin operon remained unaffected during HS, the tungsten transport protein WtpA was substantially downregulated under HS and CS conditions. Comparing the expression levels of the control condition for the regulation group, we observed that HS triggered extensive reprogramming of the transcriptome (Figure 2 H). Initially low transcribed genes are highly upregulated, while genes highly transcribed during exponential growth are massively downregulated (Fig. S3). This pattern was not observed in the CS response, where only a smaller proportion of highly upregulated genes had lower expression values in the starting condition.

The results validate functional aspects of HS regulation and reveal some overlap with the CS response, warranting further investigation into the temporal dynamics of the thermal stress responses.

### Temporal dynamics of the thermal stress response at the transcriptome level

We examined the number of up- and downregulated genes across all conditions, including HS 1, HS 2, and the recovery sample, using the previously established significance and fold change cutoffs (Figure 3 A). Approximately half of the strongly induced genes remained highly upregulated, while the other half returned to unchanged or mildly induced levels compared to the control sample. Minimal to no overlap with downregulated transcripts was observed, confirming the specificity of the response. This pattern also applied to normally upregulated genes, which returned to initial levels. A similar pattern emerged for downregulated genes, although a higher proportion of initially downregulated genes became upregulated later. Samples taken 5 min after switching from HS or CS to recovery exhibited the lowest proportion of gene expression change, with only 4% being strongly up- or downregulated.

**Figure 3.**
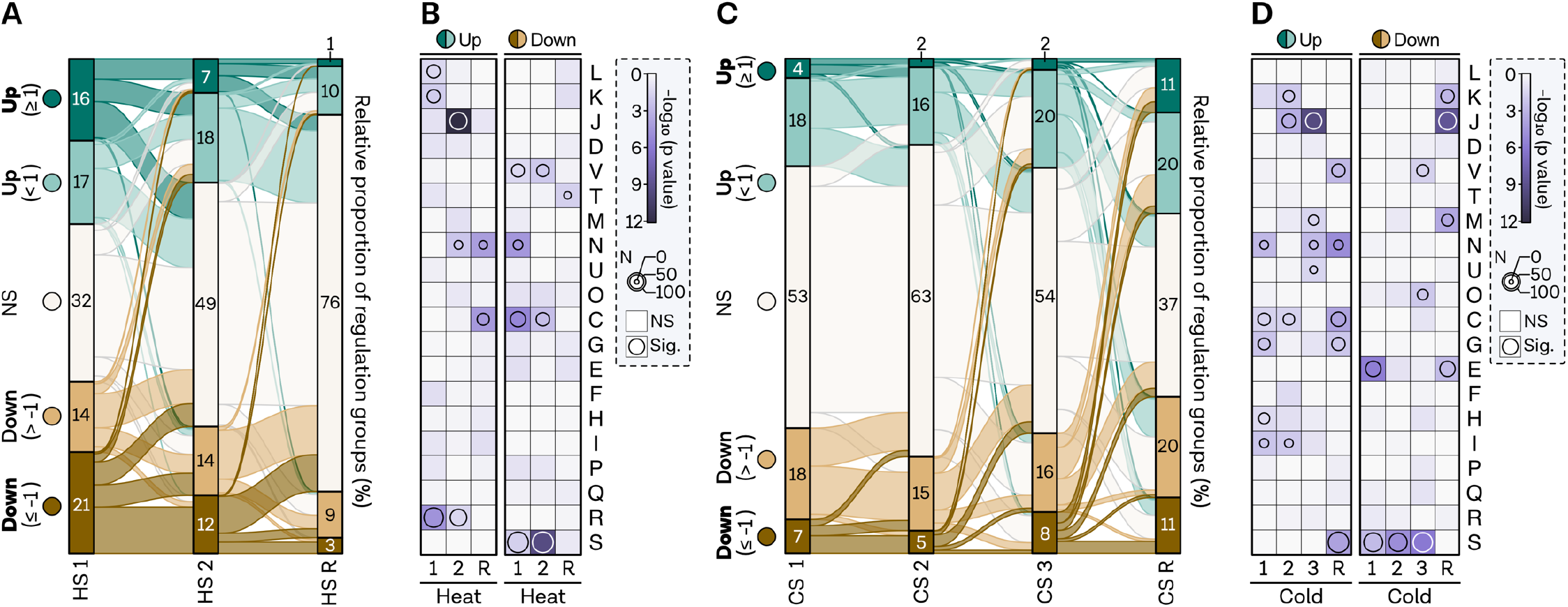
Temporal analysis shows rapid rebalancing of transcriptome changes following HS, while CS elicits multiple responses. **A**, Flow diagram visualizing the total number and interconnected genes between conditions (x-axis) in the HS experiment for each regulatory group: strongly upregulated (dark green, bold font), upregulated (light green), non-regulated (white), strongly downregulated (dark brown, bold font), and downregulated (light brown). The flow diagram size is plotted to scale. **B**, Gene set enrichment analysis of archaeal clusters of orthologous genes (arCOGs) for all three HS conditions. Significance levels are indicated by a continuous color bar from white to dark pink. Genes with a p-value < 0.05 are considered significantly overrepresented and highlighted by a circle, with circle size reflecting the number of genes in the category. For better comparison, up and down categories include all upregulated and downregulated genes, respectively, regardless of fold changes. **C**, Flow diagram for the CS experiment displaying the total number and interconnected genes between conditions (x-axis) for each regulatory group, as described in panel A. The flow diagram size is plotted to scale. **D**, Gene set enrichment analysis of arCOGs for all four CS conditions, with significance levels indicated by a continuous color bar from white to dark pink. Genes with a p-value < 0.05 are considered significantly overrepresented and highlighted by a circle, with circle size reflecting the number of genes in the category. For better comparison, up and down categories include all upregulated and downregulated genes, respectively, regardless of fold changes

Gene enrichment analysis revealed a counter-regulation indicating a rapid recovery response for some cellular processes. Genes with transcription and replication annotations were overrepresented in the upregulated groups, but this pattern was not observed in the prolonged HS 2 condition, suggesting that extended heat exposure elicits a distinct response (Figure 3 B). Instead, translation-related genes are overrepresented after extended heat exposure, indicating a switch from protection to maintaining translation. Additionally, cell motility (category N) was induced and remained enriched even after recovery, alongside energy production, which is overrepresented in downregulated genes at HS 1 and HS 2. Defence mechanisms (category V) were shut down during the immediate HS response. Notably, many currently uncharacterized genes were either shut down or upregulated during HS 1 and HS 2.

In the CS response, the number of highly up- or downregulated genes remained consistently low across all CS conditions (Figure 3 C). CS 1 and CS 2 exhibited a similar time-dynamic pattern as HS, with minimal to no overlap between up- and downregulated genes. In contrast, the 24-hour response (CS 3) showed slightly more overlap between groups, with some genes that are not differently expressed after 2 hours (CS 2) but altered after 24 hours. However, a set of highly upregulated genes remained upregulated. Interestingly, the recovery condition displayed the most significant changes. As we hypothesized that this condition could induce a heat-shock-like reaction, we compared the results to a study using an HS setup from 90 to 97°, finding the best correlation to the CS recovery from all of the conditions (Fig. S4) (49).

To further investigate this, we conducted functional enrichment analysis. Energy-related categories (C and G) were overrepresented in almost all upregulated CS conditions, including recovery. Translation-related genes exhibited a delayed response in both HS and CS but were immediately counter-regulated in the CS recovery. Transcription was overrepresented in upregulated genes earlier than translation. In contrast, most downregulated genes in the CS response have not yet been described. Only amino acid transport appears to be silenced, while the rest of the response is nonspecific.

In summary, differential gene expression and functional enrichment analysis of the RNA-seq data indicate that HS triggers substantial transcriptome changes, which is balanced remarkably quickly. In contrast, CS induces multiple responses, especially after a prolonged incubation time.

### Investigating basal features of the archaeal transcriptome landscape in HS and CS regulation

Next, we investigated whether stress-induced genes share similar regulatory sequence features with genes under normal expression, highlighting their potential role in essential cellular functions.

We first examined promoter elements, which have been shown to contribute to gene expression, although escape of the transcription elongation complex has been found to be the rate-limiting step during exponential growth (54). In *P. furiosus*, genes with archaea-typical BRE-TATA promoters display higher expression levels during exponential growth (Figure 4 A,B) (47). To determine whether stress-regulated genes are equipped with strong promoters, we analyzed the promoter strength of HS 1 and CS 1 induced genes, finding no difference compared to non-differential expressed genes (Figure 4 C). Notably, highly upregulated genes under CS conditions are all leadered, with a 5’ UTR of at least nine nucleotides possibly allowing efficient loading of the ribosome (Figure 4 D). Accordingly, more genes downregulated during CS are leaderless, which is significant in CS 1, CS 2, and the recovery condition (Figure 4 E).

**Figure 4.**
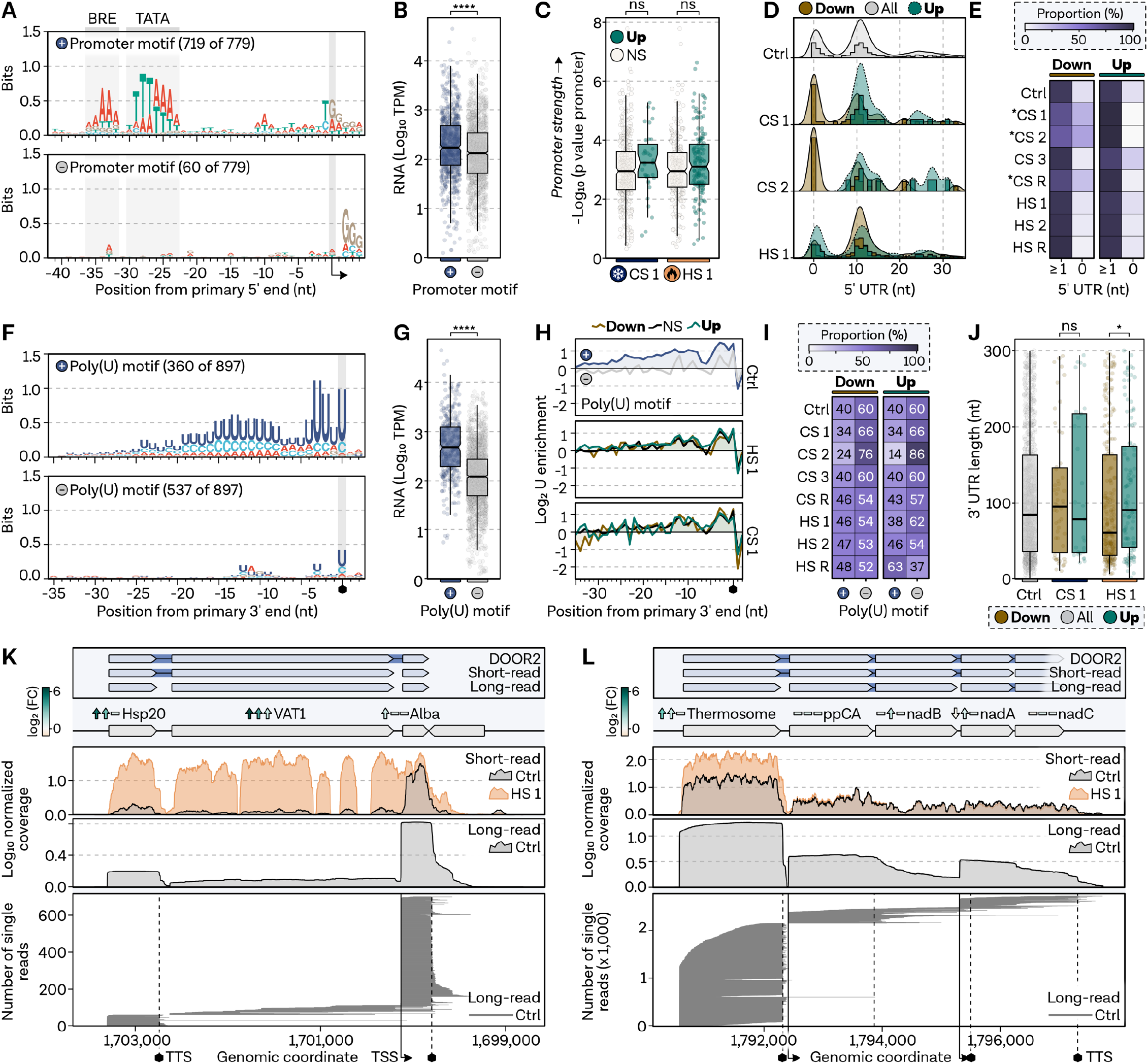
Proposed Stress-induced genes are equipped with archaea-typical regulatory sequences features. **A**, Position-specific promoter motifs of primary transcripts from (47), grouped by MEME motif search into classical archaeal promoter motif or not, based on the presence of BRE and TATA elements. **B**, Transcript abundance (TPM-normalized counts under control conditions) for genes with a promoter motif (blue) and without (grey). Box edges delineate 1st and 3rd quartiles, center line represents median, and whiskers indicate points within 1.5x interquartile range. Welch’s t-test used to assess differences between groups; significance level **** indicates p-values < 0.0001. **C**, Promoter strength comparison, estimated by p-values of MEME-detected promoter motifs (mode: one occurrence per site), for highly upregulated (dark green) and unchanged (white) genes under CS 1 and HS 1. Box plot parameters as in B; ns signifies p-values >= 0.05. **D**, 5’ UTR length comparison for highly upregulated or downregulated genes against total distribution (Ctrl, 779). Density plots shown with bars in a window size of 1 and overlaid densities. **E**, Proportion of leaderless (5’ UTR = 0) and leadered (5’ UTR >= 1) transcripts in highly upregulated and downregulated groups across all conditions. Chi-square test compares each stress condition to Ctrl sample distribution. Relative proportion color-coded from white (0) to dark purple (100%); significance (p < 0.05) indicated by an asterisk. **F**, Position-specific terminator motif based on primary 3’ ends derived from Term-seq experiments, grouped by MEME search for genes with poly(U) signal (blue) or not (white). **G**, Transcript abundance (TPM-normalized counts under control conditions) for genes with terminator poly(U) motif (blue) and without (grey). Box plot parameters as in B; significance level **** indicates p-values < 0.0001. **H**, Nucleotide enrichment meta-analysis compares nucleotide content of each group in a position-specific manner to randomly selected intergenic positions (n = 100,000). Log_2_ enrichment shown for all primary 3’ ends with poly(U) motif (blue) or not (grey), and highly upregulated and downregulated genes in HS 1 and CS 1. **I**, Proportion comparison of genes in regulatory groups with poly(U) signal or not, for all conditions. Note that an insufficient number of genes was detected for the CS 2 sample for robust statistical testing. Chi-square test checks whether the proportion of genes with poly(U) terminator differs significantly from Ctrl condition; no significant difference at p < 0.05. **J**, 3’ UTR comparison of regulatory groups to the Ctrl condition. Box plot parameters as in B; significance levels of * and ns indicate p-values < 0.05 and >= 0.05, respectively. **K**, Operon analysis using long-read PCR-cDNA Nanopore sequencing for HSP20 operon and **L**, thermosome operon. Top panel: annotated operons from DOOR2 database, previous short-read RNA-sequencing, and long-read Ctrl condition sequencing. Genes visualized as rectangles, strand indicated by arrow direction, and operons connected by blue background lines. Next panel: current gene annotation and log_2_ fold changes on RNA level for three HS conditions (1, 2, R) from left to right. Significance indicated by up or down arrows. Profiles of mean normalized coverage values from Nanopore and short-read RNA-sequencing, color-coded as Ctrl (grey) and HS 1 (orange). The last panel presents a single-read analysis of Nanopore reads. Each line represents one full-length sequenced read, sorted by their read start position. Vertical lines indicate primary start sites identified in (47), while dashed lines represent primary 3’ ends from short-read Term-sequencing.

Next, we employed Term-seq to survey the termination landscape of the *P. furiosus* transcriptome, identifying 897 enriched 3’ ends downstream of protein-coding transcripts under exponential growth conditions (Table S2). In agreement with previous findings for archaea, we discovered a long poly(U)-stretch of approximately 16 bases enriched at the 3’ end (Figure 4 F) (55, 56). Although many genes (538 of 897) lack a robust poly(U)-signal, termination generally does not appear to be triggered by secondary structures (Fig. S5). Genes exhibiting a strong poly(U)-motif at the 3’ end are highly expressed under optimal exponential growth conditions (Figure 4 G). Assessment of poly(U) enrichment in highly up- and downregulated genes under HS and CS conditions revealed no difference compared to the control sample, suggesting that numerous stress genes are terminated by poly(U)-signals and therefore equipped for efficient termination (Figure 4 H,I). Additionally, no difference in 3’ UTRs was observed between up- or downregulated genes in any condition except HS 1, where downregulated genes exhibit significantly shorter 3’ UTRs (Figure 4 J).

Many genes associated with HS and CS in *P. furiosus* are predicted or validated to be transcribed within an operon, impacting transcriptional and translational regulation (47). Upon investigation of the operon organization of certain signature HS response genes, we found that the differential gene expression data did not correspond with operonization (Figure 4 K,L). To better understand the transcription of these genes, we collected long-read RNA-seq data using Nanopore sequencing of PCR-amplified cDNAs from the control condition. We specifically investigated an operon containing HSP20, VAT1 and the highly abundant DNA-binding protein AlbA. Single-read analysis confirmed 3’ end data gathered from Term-seq and primary 5’ end positions, suggesting a primary stop immediately after HSP20 and separate transcription start sites for each gene (Figure 4K). This is consistent with Phr regulation of HSP20 and VAT1, as binding sites precede each gene. Interestingly, the stress condition HS 1 explains why the algorithm used in the 2019 paper, based on short-read data analysis of mixed RNA conditions, identified both of these genes in an operon (47). This observation is further supported when examining the thermosome operon, which, according to the long-read data, is distinctly transcribed as a single gene with very minimal readthrough rather than alongside other genes as annotated in the DOOR2 database (Figure 4L).

### Integrated transcriptomics and proteomics unveil shared functional responses and moderate correlation between total RNA and protein expression values

To further elucidate the molecular mechanisms underlying thermal stress response, we next analyzed the proteomics data (quantities and iBAQ values listed in Table S3). Focusing on the HS response, where we observed a distinct response in the PCA (Figure 1 G), we identified 23% of the genes as upregulated and 26% as downregulated (Figure 5 A). Notably, log_2_-fold changes at the protein level were smaller as compared to the RNA level, prompting us to forgo applying an additional threshold for group description to ensure robust analysis. Key HS response proteins, such as the Myo-Synthase and Hsp20, displayed significant upregulation. Time-dependent analysis revealed no overlap between up- and downregulated groups during HS, a trend more pronounced than in the transcriptomics data. Moreover, a larger number of proteins remained affected during the recovery condition compared to the RNA-seq data, indicating that protein-level reactions occur at a slower pace (Figure 5 B). For CS, we observed that 16% of proteins were upregulated after 20 minutes and 18% were downregulated, with virtually no change between the control condition and the 2-hour protein sample (Figure 5 C). After 24 hours, substantial alterations in the proteome were evident, with regulation rapidly reverting in the recovery sample.

**Figure 5.**
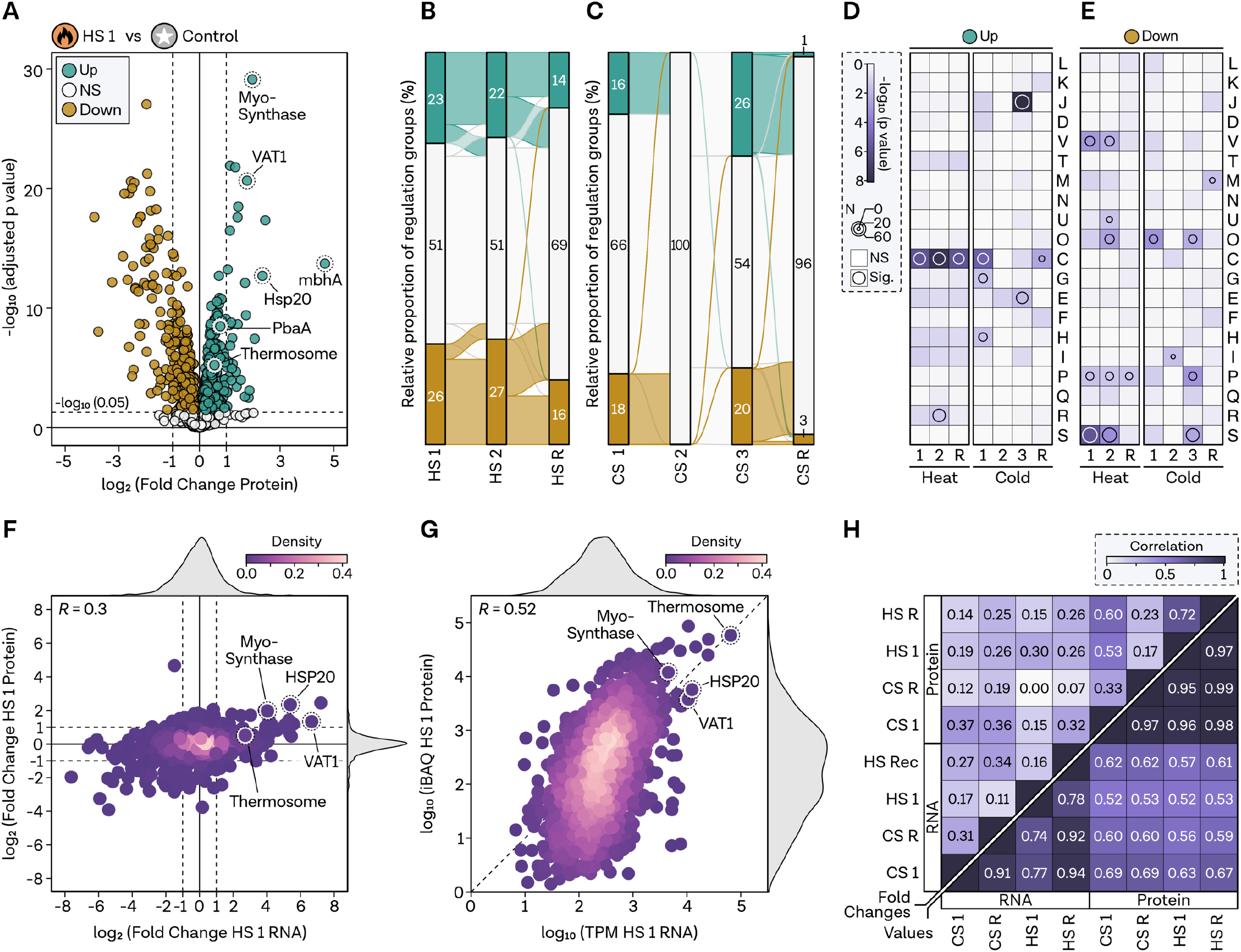
Proteomics count values but not fold changes are moderatly correlated to RNA levels. **A**, Volcano plot displaying log_2_ fold changes in protein levels (x-axis) and −log10 (adjusted p-value, padj) on the y-axis comparing HS 1 (5 minutes) with control condition. Protein-coding genes are categorized based on significance and fold changes, color-coded as upregulated (padj < 0.05, log_2_FC > 0, green), non-regulated (NS, padj >= 0.05, white), and downregulated (padj < 0.05 and log_2_FC < 0, brown). HS genes are highlighted. Genes were assessed at a significance level of 0.05, indicated by a dashed line in the plot. **B**, HS and **C**, CS flow diagram visualizing the total number and interconnected genes between conditions on the x-axis for each regulatory group, as described in panel A. **D**, Gene set enrichment analysis of archaeal clusters of orthologous genes (arCOGs) for all three HS and **E**, all four CS conditions. Significance level is indicated by a continuous color bar from white to dark pink. Genes with a p-value < 0.05 are considered significantly overrepresented in the category and are highlighted by a circle. The circle size reflects the number of genes in the category. **F**, Comparison of log_2_ fold changes and **G**, normalized TPM and iBAQ values of HS 1 measured at RNA (x-axis) and protein (y-axis) levels. HS genes are highlighted. Pearsońs correlation coefficient is shown in the top left. Density of plotting is color-coded. **H**, Correlation matrix (Pearsońs correlation coefficient) of pairwise comparisons based on fold changes (upper left corner) and normalized count values (bottom right).

Functional enrichment analysis demonstrated that overrepresented groups exhibited greater consistency compared to the RNA sequencing data (Figure 5 D). Energy-related genes were overrepresented across all upregulated HS categories, as well as in the CS 1 and recovery conditions, suggesting potential similarities at the protein level. Additionally, after 24 hours of CS, translation-related genes were overrepresented, which had already been observed at the RNA level. Downregulated genes under HS included defence mechanisms, ion transport, and, in part, post-transcriptional modifications and chaperones. Numerous currently uncharacterized proteins were also downregulated under HS (Figure 5 E).

Integrating our findings with the transcriptomics data, we identified a low correlation between fold changes; however, the same trend was evident for previously known HS-responsive genes (Figure 5 F). The correlation improved but remained moderate when comparing total expression values, normalized by either TPM for transcripts or iBAQ value for proteins (Figure 5 G). Upon comparing various conditions, we determined that the correlation between CS conditions was generally higher, particularly at the count level, where a notable correlation of 0.69 emerged between RNA and protein iBAQ values in CS 1 (Figure 5 H).

### Identifying signature genes and common principles for HS and CS response

Analyzing the response regulated by the TF Phr in *P. furiosus*, we observed a time-dependent trend in RNA and protein values relative to the control sample, indicating a rapid transcriptome-level response followed by a more prolonged, less dynamic response at the protein level (Figure 6 A, B). To further examine regulatory aspects, we conducted a cluster analysis to identify groups sharing temporal trends in transcript and protein levels (Summary of all data listed in Table S4). Therefore, we performed hierarchical clustering of z-score normalized log_2_-fold changes, accounting for the varying sensitivities of RNA-seq and MS (Figure 6 C). In the HS response, we identified distinct clusters displaying either similar (cluster 3, 4) or contrasting trends in RNA and protein levels, particularly during HS 1 (cluster 1, 2) and HS 2 (cluster 5). Log_2_-fold change analysis and arCOG enrichment of these clusters facilitated the integrative analysis in a functional context (Figure 6 D,E,F).

**Figure 6.**
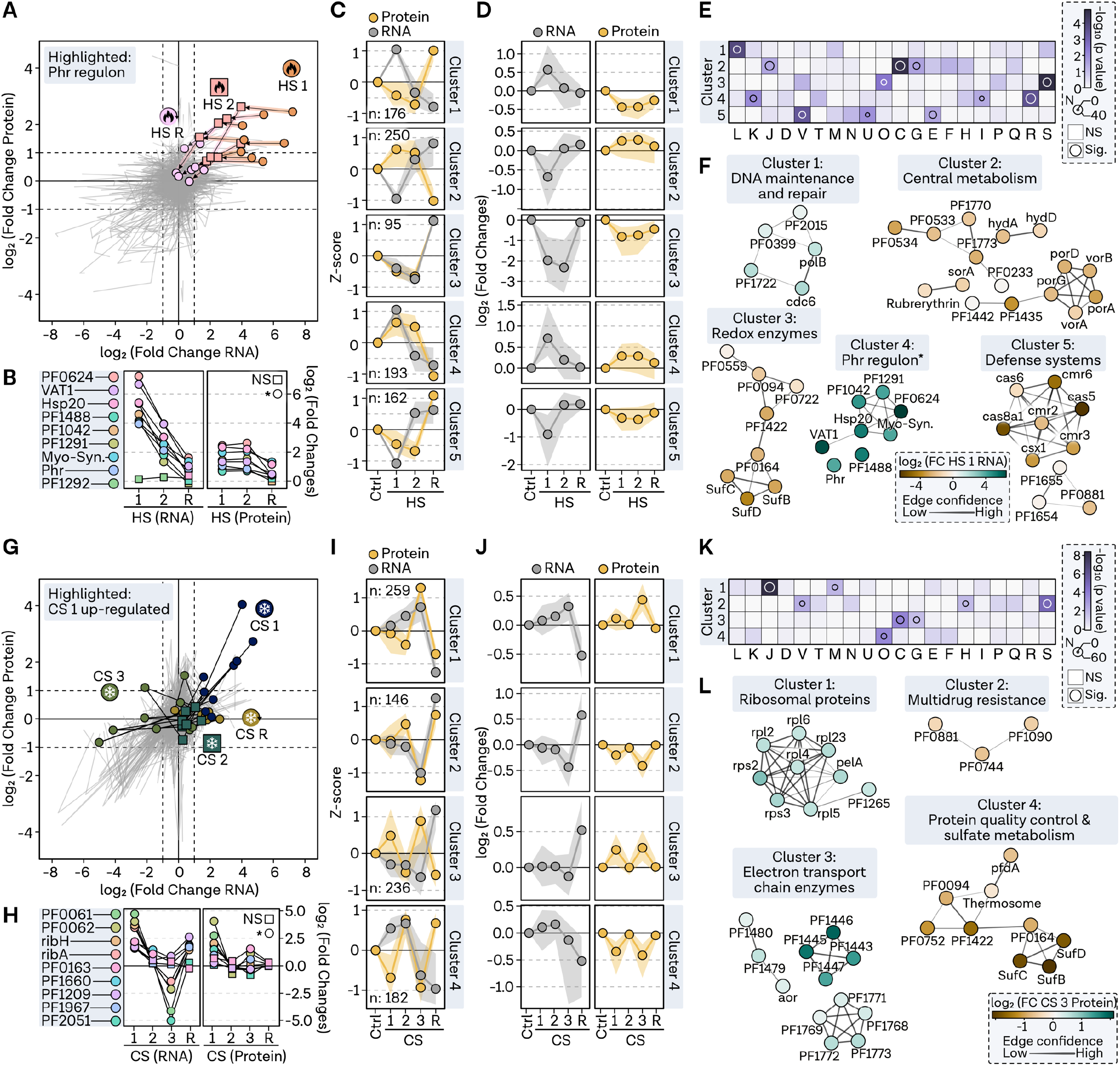
Cluster analysis identifies signature genes in HS and CS responses. **A**, Pathway plot illustrating log_2_ fold changes at RNA level (x-axis) and protein level (y- axis) in a line plot from condition HS 1 via HS 2 to recovery condition. Genes of the Phr regulon are highlighted with colors and points. **B**, Fold changes for genes presented in panel A. Significant regulation in respective conditions is indicated by a circle (padj < 0.05) or a rectangle. **C**, Comparison of median z-score values shown as a point in each cluster (row) and condition (column) for protein (yellow) and RNA (grey) values. Shaded area represents the interquartile range. **D**, Log_2_ fold changes of genes sorted by clusters. Points indicate median, while shaded area shows interquartile range. **E**, Gene set enrichment analysis of archaeal clusters of orthologous genes (arCOGs) for five selected clusters. Significance level is represented by a continuous color bar from white to dark pink. Genes with a p-value < 0.05 are considered significantly overrepresented in the category and are highlighted by a circle. Circle size reflects the number of genes in the category. **F**, Functional enrichment analysis of genes from each cluster was performed by selecting all significantly regulated genes in the most significantly overrepresented arCOG category. Confidence in predicted interactions, according to the STRING database, is indicated by line thickness. Only genes with at least one connection are shown. Genes are depicted as points and colored based on the fold change at RNA level in condition HS 1. **G**, Pathway plot illustrating log_2_ fold changes at RNA level (x-axis) and protein level (y-axis) in a line plot from condition CS 1, through CS 2 and CS 3 to recovery condition. The nine genes with the highest upregulation at RNA level (CS 1) are highlighted with colors and points. **H**, Fold changes for genes presented in panel G. Significant regulation in respective conditions is indicated by a circle (padj < 0.05) or a rectangle. **I**, Comparison of median z-score values shown as a point in each cluster (row) and condition (column) for protein (yellow) and RNA (grey) values. Shaded area represents the interquartile range. **J**, Log_2_ fold changes of genes sorted by clusters. Points indicate median, while shaded area shows interquartile range. **K**, Gene set enrichment analysis of archaeal clusters of orthologous genes (arCOGs) for four selected clusters. Significance level is represented by a continuous color bar from white to dark pink. Genes with a p-value < 0.05 are considered significantly overrepresented in the category and are highlighted by a circle. Circle size reflects the number of genes in the category. **L**, Functional enrichment analysis of genes from each cluster was performed by selecting all significantly regulated genes in the most significantly overrepresented arCOG category. Confidence in predicted interactions, according to the STRING database, is indicated by line thickness. Only genes with at least one connection are shown. Genes are depicted as points and colored based on the fold change at protein level in condition CS 3.

HS cluster 1 featured DNA maintenance and repair genes upregulated at the RNA level but downregulated at the protein level during HS 1, such as the cell cycle regulator cdc6, the DNA polymerase subunit B, the histone A2 (PF1722), and hef-associated nucleases PF0399 and PF2015. Conversely, HS cluster 2 displayed counter-regulation, with upregulation of protein levels and downregulation of RNA levels. This cluster was significantly enriched in genes associated with central metabolism and translation-related processes. HS cluster 3 exhibited striking consistency in regulation, with genes related to post-translational modification, protein turnover, and chaperones initially downregulated during the immediate shock and subsequently recovering. This pattern was particularly evident for redox enzymes like SufB, SufC, and SufD. HS cluster 4 displayed the regulation expected for a HS protective response and is characterized by genes rapidly upregulated under HS 1 at the RNA level and a prolonged upregulating effect on the protein level. In this case the signature genes were not selected based on arCOGs but on experimentally identified Phr targets, all of which were present in this cluster. However, while some transcription-related genes are overrepresented in cluster 4, we did not find some of the genes initially thought to contribute to thermal fitness in the cluster. One prominent example are nucleoid-associated proteins (NAPs), like AlbA (PF1881), TrmBL2 (PF0496) and histones A1 (PF1831) and A2 (PF1722), that are not uniformly regulated (Fig. S6 A). Nevertheless, we could confirm that *P. furiosus* has a high investment of the total protein to the NAPs, especially histone A2 and AlbA with protein levels increasing up to >5% under HS conditions (Fig. S6 B) (17). Lastly, cluster 5 encompassed defense systems downregulated at the RNA level during HS 1, with protein levels remaining down for an extended period, displaying an opposite trend to cluster 4. This cluster contained genes related to defense systems transcribed from the Cas locus 1, including Cmr (Type III-B) and Cst (Type I-G), as well as transporter-related genes. The refined clustering and functional analysis provided a clear depiction of the various regulatory patterns observed in the HS response of *P. furiosus*, highlighting the complexity of the molecular mechanisms underlying this process.

Re-evaluating the genes in HS cluster 4, we hypothesized that the Phr regulon may encompass additional targets not yet identified experimentally. Indeed, through motif analysis and comparison to the RegPrecise database, which houses transcriptional regulons based on comparative genomics, we discovered additional targets with palindromic Phr binding motifs, expanding the initial 10-gene group (Fig. S7) (57). We identified the proteasome-assembling chaperone PbaA and its complex partner PF0014, the predicted transcriptional regulator PF1932, and the KaiC domain-containing protein PF1931 as additional targets strongly upregulated under HS (Fig. S7 A,B). By comparing the Phr motif with the upstream sequences of predicted targets, we noted that genes predicted to have a motif but not exhibiting the typical Phr-mediated response at the RNA and protein levels, such as PF0239 and PF1117, deviate from the consensus sequence (Fig. S7 C). Additionally, we confirmed previous analyses showing that PF0321 has two start sites, with only the more distant site being under the control of the Phr motif, resulting in a weaker response.

In analyzing the regulatory dynamics during CS response, we observed distinct short- and long-term responses, which were further investigated. Interestingly, the riboflavin operon and several highly upregulated genes during CS 1 were also affected at the protein level but were strongly downregulated during CS 1, while protein levels remained unchanged or even increased (Figure 6 G,H). Performing cluster analysis similar to the HS samples, we identified distinct clusters that are discussed in the following (Figure 6 I): CS cluster 1 featured proteins especially upregulated at CS 3, with increasing upregulation at the RNA level over time, encompassing translation-related and membrane proteins (Figure 6 J,K). Notably, ribosomal proteins were upregulated at the protein level after 24 hours, counter-regulated after recovery at 95°C. Interestingly, PF1265, a tRNA/rRNA cytosine-C5 methylase, was identified in this cluster. Further analysis of potential tRNA/rRNA-modifying genes revealed that PF1265 was indeed the only gene upregulated under CS 3 at both RNA and protein levels (Fig. S8). Conversely, KsgA expression was downregulated at the protein level after HS, and only the Fmu homologue PF0666 and the tRNA methyltransferase PF1871 were upregulated at both levels during HS. CS cluster 2 is characterized by coordinated downregulation at both RNA and protein levels, particularly after prolonged CS. This cluster contains several unknown genes, as well as transporters related to multidrug resistance. CS cluster 3 exhibits upregulation at CS 1 and CS 3 time points on the protein level and is enriched in metabolism-related genes, including electron transport chain enzymes. In contrast, CS cluster 4 displays downregulation at the protein level immediately following CS. Interestingly, this cluster contains proteins involved in protein quality control, such as the thermosome and prefoldin alpha, and Fe-S cluster assembly proteins SufB, SufC, and SufD, which are implicated in sulfate metabolism.

Considering that some of these genes are regulated by the TF SurR (PF0095), we investigated the overlap between the thermal stress response to other known TF-regulons in *P. furiosus*. Therefore, we examined experimentally verified regulons, such as SurR, CopR (PF0739), TrmBL1 (PF0124), TFB-RF1 (PF1088), as well as predicted regulons like cobalamin biosynthesis regulation by CblrR (unknown), riboflavin biosynthesis operon regulation by RbkR (PF0988), and thiamin transport regulation by ThiR (PF0601) (Fig. S9) (40, 57–61). Many genes controlled by validated or yet unknown transcription factors exhibit significant regulation at both the RNA and protein levels, suggesting either TF regulatory networks that affect one another or secondary effects. While genes upregulated by SurR during the primary S^0^ response also exhibit upregulation under CS and HS, genes downregulated by SurR display consistent downregulation under the tested conditions at the RNA level. Intriguingly, targets of the copper-regulator CopR are substantially downregulated, especially genes involved in binding and transporting metal ions, such as PF0723.

To explore additional possible determinants of RNA and protein fate, we analyzed sequence features, including the codon adaptation index (CAI) and GC content of coding regions (Summary of all data listed in Table S4) (Fig. S10). We found that all HS clusters characterized by upregulation of the proteome (clusters 2 and 4) exhibit significantly higher CAI values compared to the overall distribution (Fig. S10 A). Cluster 2 is also characterized by genes with higher GC content, while cluster 1 has lower than expected GC values (Fig. S10 B). Regarding the CS clusters, not only do the clusters characterized by protein upregulation exhibit higher CAI values, but also cluster 4. However, the GC content is only significantly higher in clusters 1 and 3 (Fig. S10 C,D).

In summary, while the response to HS is regulated by Phr and possibly encoded at the sequence level, other effects are likely controlled by currently undetermined features, as well as post-transcriptional and post-translational regulatory mechanisms.

## Discussion

In this study, we investigated the response of the hyperthermophilic archaeon *P. furiosus* to HS and CS, unveiling complex patterns of gene expression and protein regulation. Our results indicate a well-coordinated, temporally-organized response, utilizing various adaptive mechanisms for distinct cellular processes and pathways.

HS triggers extensive reprogramming of the transcriptome, upregulating numerous heat shock proteins that serve to prevent cellular damage. We confirmed HS-induced upregulation of predicted targets of the transcriptional regulator Phr on both RNA and protein levels in a time-resolved manner, thus broadening its known regulon (40). Moreover, we demonstrated that stress-induced genes generally possess archaea-typical promoter and transcription terminator sequences. This suggests that these genes are silenced under normal growth conditions but can be rapidly activated, as observed for Phr-regulated genes. While HS primarily leads to the upregulation of proteins related to energy production and transcription-related processes, we observed the downregulation of defence mechanisms, including the CRISPR-Cas system. Silencing of these systems during stress might be attributed to energy conservation or preventing inadvertent activation of the immune response, which could adversely affect genome stability, especially since RNA and DNA targeting CRISPR-Cas systems coexist in *Pyrococcus* (62, 63).

While we and others found a number of proteins that confer a specific heat shock response, we did not identify proteins that confer protection against CS. The CS response appears to be characterized by two distinct phases: an initial phase focused on energy provision, followed by a phase aimed at sustaining translation. This coordinated response is evident at both RNA and protein levels and rapidly reverts during recovery. Regulation during CS is more subtle compared to HS, with multiple responses observed at both short and longer-term CS on RNA and protein levels. While previous studies investigating the CS response in (hyper-)thermophilic archaea have identified some cold-induced genes, only a subset of these are reflected in our data. This discrepancy might be explained by the experimental conditions used in earlier studies, which primarily involved temperatures at the lower growth limit as the CS temperature (52, 64, 65). Nevertheless, a 4°C shock is a plausible scenario for these organisms, considering their exposure to surrounding cold seawater that may also penetrate the porous material of black smokers (66).

Interestingly, it has been reported that general adaptations of psychrophilic archaea share some characteristics with CS response in hyperthermophiles (65). One of these signature domains found in psychrophiles is TRAM, which is universally distributed and functions via RNA chaperone activity (67–69). The protein containing this domain, possibly mimicking the bacterial cold shock protein A function in archaea, is also present in *P. furiosus* (PF1062). While it is downregulated under early CS conditions, it is significantly upregulated after 24 h on the protein level. In addition to the investigation of single cold-induced genes, a global quantitative proteomics study in the cold-adapted *Methanococcoides burtonii* demonstrated that the abundance of ribosomal subunits peaked at 4°C (70). Our analysis supports this observation, as we detected a time-dependent upregulation of ribosomal proteins. Interestingly, this may be connected to the finding that genes upregulated during early CS in our study consistently possess a 5’ leader sequence, compared to approximately 15% leaderless genes observed under normal conditions. The presence of a 5’ leader sequence, including a Shine-Dalgarno site, could potentially facilitate ribosome recruitment and translation initiation factor binding, thereby ensuring efficient translation of cold-responsive genes under these challenging conditions (71). Moreover, this arrangement may allow for more intricate post-transcriptional regulatory mechanisms, such as RNA-binding protein interactions or secondary structure formation, which could enable fine-tuning of gene expression. However, the mechanism of leaderless translation remains unclear, complicating functional comparisons (72–74).

The overlap of transcription regulons examined in this study underscores the complex cellular responses to environmental stressors, which may be coordinated across a broad range of conditions. Notable examples include the RbkR-regulated riboflavin (vitamin B2) operon, producing essential cofactor precursors flavin mononucleotide (FMN) and flavin adenine dinucleotide (FAD), and the ThiR-regulated thiamine (vitamin B1) operon, precursor of thiamine pyrophosphate (TPP) (53, 75, 76). Given the substantial upregulation of energy-related enzymes during CS, there may be an increased demand for maintaining proper functioning of metabolic pathways, which could explain the upregulation of these operons (77). Alternatively, secondary effects may also contribute to the observed transcriptional regulation. For example, disrupted redox homeostasis might explain the regulation of SurR targets, which is a redox-active transcriptional regulator (58, 78, 79).

Although our study has yielded significant findings, we must acknowledge certain limitations. Our experimental design focused on specific time points, potentially missing the full dynamics of gene expression changes during stress response. Time-course experiments for certain targets would offer a more comprehensive view of temporal changes in gene expression and could reveal additional regulatory mechanisms. Additionally, our experimental setup allows only for relative quantitative comparisons of the final amounts of transcripts and proteins in the cell. As such, we cannot make definitive statements regarding the neo-synthesis of RNA and proteins, which should be considered for interpretation. Furthermore, we cannot address the heterogeneity of responses on the genome and transcriptome levels. While single-cell analysis has provided deeper insights into individual cellular mechanisms during thermal stress and various growth conditions in bacteria, its application to archaea remains unexplored (80, 81). Future utilization of single-cell technologies will help uncover rare cellular states and genome plasticity of individual *Pyrococcus furiosus* cells, particularly during HS, shedding light on the diverse strategies these organisms employ to cope with extreme environments.

In conclusion, our study comprehensively analyses of the transcriptomic and proteomic responses to thermal stress in the hyperthermophilic archaeon *Pyrococcus furiosus*. We identified distinct expression patterns and regulatory mechanisms, providing valuable insights into the dynamic response mechanisms to environmental fluctuations and their control by transcription factor networks in archaea. Our findings enhance our understanding of *P. furiosus*’ remarkable adaptability and expand our knowledge of life in extreme environments.

## Material & Methods

### Growth conditions and sample preparations

*Pyrococcus furiosus* DSM 3638 was cultured in 120 ml serum bottles following the protocol previously described under anaerobic conditions at one bar excess of nitrogen (47). Cells were grown in 40 ml SME medium supplemented with 40 mM pyruvate, 0.1 % peptone and 0.1 % yeast extract for 15 hours at 95°C until they reached the mid-exponential growth phase with a cell density of 5 x 10^7^ cells/ml (47).

For cold shock treatment, cells in the mid-exponential growth phase were subjected to cold shock by rapidly cooling the medium through a 2.5 m long, 2 mm diameter viton hose flushed with anaerobic NaCl solution into a new bottle containing 1 bar excess of nitrogen (Fig. S1 B,C). The transfer took 90 seconds, and the time to reach 4 °C in an ice water bath was 160 seconds in total, which marked the beginning of the cold shock treatment. For recovery, the ice bath was replaced with a 95°C hot water bath, and a recovery sample was collected 5 minutes after the target temperature of 95°C was achieved.

For heat shock treatment, serum bottles with cells in the mid-exponential growth phase were placed in a 105°C incubator. It took 26 minutes for the cultures to reach the target temperature of 105°C, which was considered the starting point for the heat shock treatment. Samples were collected after 5 and 15 minutes of incubation at this temperature. For recovery, cultures were transferred to another incubator and allowed to cool down to 95°C, which took 10.5 minutes (Fig. S1 D).

Cells from all conditions were harvested by centrifugation at 14,000 x g for 10 minutes at 4°C and resuspended in 2 ml buffer (25 mM Tris/HCl pH 7.6, 100 mM NaCl). Samples were then divided for RNA (1.5 ml) and protein analysis (0.5 ml), followed by centrifugation at 17,700 x g for 5 minutes at 4°C. The resulting pellets were stored at −80°C until further analysis.

### RNA extraction

Total RNA isolation was performed using cell pellets, which were initially lysed by adding 1 ml of TRI Reagent (Zymo Research) and subsequently processed with the Direct-zol RNA Miniprep Plus kit (Zymo Research) following the manufacturer’s instructions. This protocol included DNase I digestion to eliminate genomic DNA contamination.

To evaluate the purity, quality, and quantity of the extracted RNA, several analytical methods were employed. Standard spectroscopic measurements were performed using a NanoDrop One spectrophotometer (Thermo Fisher Scientific). Fluorometric quantification was carried out with the Qubit RNA assay (Thermo Fisher Scientific). Lastly, RNA integrity values were assessed using a Bioanalyzer (Agilent Technologies) to ensure the suitability of the RNA samples for downstream applications.

### RNA sequencing for differential gene expression analysis

#### Library preparation and sequencing

Four independent biological replicates were prepared for each condition and subjected to differential gene expression analysis. Prior to library preparation, ribosomal RNAs were depleted from 1 µg input RNA using the Pan-Archaea riboPOOL (siTOOLs) according to the manufacturer’s instructions. Library preparation and RNA-seq were carried out as described in the Illumina “Stranded mRNA Prep Ligation” Reference Guide, the Illumina NextSeq 2000 Sequencing System Guide (Illumina, Inc., San Diego, CA, USA), and the KAPA Library Quantification Kit - Illumina/ABI Prism (Roche Sequencing Solutions, Inc., Pleasanton, CA, USA). In brief, omitting the initial mRNA purification step with oligo(dT) magnetic beads, around 5 ng of rRNA depleted archaeal RNA was fragmented to an average insert size of 200-400 bases using divalent cations under elevated temperature (94°C for 8 minutes). Next, the cleaved RNA fragments were reverse transcribed into first strand complementary DNA (cDNA) using reverse transcriptase and random hexamer primers. Actinomycin D was added to allow RNA-dependent synthesis and to improve strand specificity by preventing spurious DNA-dependent synthesis. Blunt-ended second strand cDNA was synthesized using DNA Polymerase I, RNase H and dUTP nucleotides. The incorporation of dUTP, in place of dTTP, quenches the second strand during the later PCR amplification, because the polymerase does not incorporate past this nucleotide. The resulting cDNA fragments were adenylated at the 3’ ends and the pre-index anchors were ligated. Finally, DNA libraries were created using a 15 cycle PCR to selectively amplify the anchor-ligated DNA fragments and to add the unique dual indexing (i7 and i5) adapters. The libraries were bead purified twice and quantified using the KAPA Library Quantification Kit. Equimolar amounts of each library were sequenced on an Illumina NextSeq 2000 instrument controlled by the NextSeq 2000 Control Software (NCS) v1.4.1.39716, using one 50 cycles P3 Flow Cell with the dual index, single-read (SR) run parameters. Image analysis and base calling were done by the Real Time Analysis Software (RTA) v3.9.25. The resulting .cbcl files were converted into .fastq files with the bcl2fastq v2.20 software.

Library preparation and sequencing were performed at the Genomics Core Facility “KFB - Center of Excellence for Fluorescent Bioanalytics” (University of Regensburg, Regensburg, Germany; www.kfb-regensburg.de).

#### Differential gene expression analysis

Raw sequencing reads in fastq format were filtered for quality and trimmed using fastp (v. 0.23.2) to remove low-quality bases and adapter sequences (--cut_front -- cut_tail -q 30) (82). Reads mapping to ribosomal RNAs were bioinformatically removed from the trimmed reads using the sortmeRNA tool (v. 4.3.6) based on sequences in the SILVA database. In a next step, the rRNA-depleted reads were aligned to the *Pyrococcus furiosus* DSM 3638 reference genome (NCBI: CP023154.1) using Bowtie2 (v. 2.5.0) with default parameters (83). The resulting sequence alignment files (SAM) were converted to binary mapping format (BAM) using samtools (84).

Differentially expressed genes from RNA-seq count data were identified following the recommendations in the Bioconductor vignette of the DESeq2 package (85). Briefly, featureCounts (RSubread package v. 2.10.5) was used to calculate the count matrix based on a custom GTF file generated by filtering the *P. furiosus* DSM 3638 GFF annotation file downloaded from the NCBI for protein-coding genes (column biotype) (86). Principal component analysis (PCA) was performed on variance stabilizing transformed (VST) data, and outlier replicates (HS 1 replicate 1, HS 2 replicate 4, HS R replicate 1, CS R replicate 2) were removed from the dataset after visual inspection. Differential expression analysis was conducted by comparing each of the cold or heat shock conditions with the control condition in a pairwise manner.

### Term-seq

*Library preparation and sequencing:* For efficient 3’ adapter ligation, 1 µg DNase-treated RNA (control condition, 95°C, mid-exponential growth phase) was mixed with 1 μl RNA 3’ adapter containing four degenerate bases (5’-amino-NNNNAGATCGGAAGAGCGTCGTGTAGGGAAAG-phosphate-3’, Microsynth, 150 μM), 2.5 μl 10x T4 RNA ligase buffer, 2.5 μl ATP (10 mM), 2 μM DMSO (100%), 9 μl PEG8000 (50%), 2.5 μl T4 RNA ligase 1 (10 U/μl, NEB), and 0.5 μl RiboLock RNase Inhibitor (40 U/μl) and filled up to a total volume of 25 μl. The reaction was incubated at 23°C shaking (450 rpm) for 2.5 h and cleaned up with 2.2 x vol AMPure XP Beads.

Subsequently, rRNAs were depleted using the Pan-Archaea riboPOOL (siTOOLs) according to the manufactureŕs instructions. 3’-ligated and rRNA-depleted RNAs were split in half and fragmented by adding 1 μl 10x RNA Fragmentation Reagent (Ambion) and incubation at 70°C for 7.5 min and 15 min, respectively. The reactions were stopped by the addition of 1 μl stopping solution (200 mM EDTA pH 8.0), pooled and cleaned up with 2.2 x vol AMPure XP Beads. For cDNA synthesis, fragmented RNAs were first mixed with 1 μl RTP primer (5’-TCTACACTCTTTCCCTACACGACGCTCTTC-3’, 10 μM) and incubated for 5 min at 65°C. Subsequently, samples were placed on ice and 4 μl 5x First-Strand Buffer, 1 μl DTT (100 mM), 2 μl dNTPs (10 mM) and 2 μl SuperScript III (200 U/μl) was added. Synthesis was performed for 60 min at 50°C after which the enzyme was heat inactivated for 15 min at 70°C. Next, RNase digestion was performed by adding 1 μl RNase H (5 U/μl) and incubating the samples for 30 min at 37°C. The reaction was cleaned using 2.2 x vol AMPure XP beads. To reduce primer dimer formation, cDNA was mixed with 1 μl 10 μM 3’ RNA adapter and denatured for 3 min at 95°C in a thermoblock. Subsequently, the samples were cooled to 23°C with an initial cooling rate of 10 sec/°C until 72°C at which the cooling rate was lowered to 15 sec/°C. Next, 2.5 μl 10x T4 RNA ligase buffer, 2.5 μl ATP (10 mM), 2 μM DMSO (100%), 9.5 μl PEG8000 (50%), 2.5 μl T4 RNA ligase 1 (10 U/μl, NEB) and 1 μl 3’ cDNA adapter (5’-amino-NNNNAGATCGGAAGAGCACACGTCTGAACTCCAGTCAC-phosphate-3’, Microsynth, 150 μM) were added to the hybridized cDNA, incubated at 23°C shaking (450 rpm) for 6 h and cleaned up with 2.2 x vol AMPure XP beads. Finally, cDNA libraries were amplified using a 12 cycle PCR with NEBNext^®^ Multiplex Oligos for Illumina protocol (NEB) and a standard Phusion polymerase protocol (NEB). Amplified libraries were cleaned up after gel electrophoresis (selected size between 200 bp to 500 bp) by a NucleoSpin Gel and PCR Clean-up kit (Macherey-Nagel) according to the manufacturer’s instructions for high percentage agarose gels.

Sequencing was performed at the Genomics Core Facility “KFB -Center of Excellence for Fluorescent Bioanalytics” (University of Regensburg, Regensburg, Germany; www.kfb-regensburg.de) and carried out as described in the Illumina NextSeq 500 System Guide (Illumina, Inc., San Diego, CA, USA), and the KAPA Library Quantification Kit -Illumina/ABI Prism (Roche Sequencing Solutions, Inc., Pleasanton, CA, USA). In brief, the libraries were quantified using the KAPA Library Quantification Kit. Equimolar amounts of each library were sequenced on a NextSeq 500 instrument controlled by the NextSeq Control Software (NCS) v2.2.0, using one 75 Cycles High Output Kits with the single index, paired-end (PE) run parameters. Image analysis and base calling were done by the Real Time Analysis Software (RTA) v2.4.11. The resulting .bcl files were demultiplexed and converted into .fastq files with the bcl2fastq v2.18 software.

The Term-seq protocol was applied to four biological replicates of cells grown to mid-exponential growth phase at 95°C.

### Identification of enriched 3’ ends

Fastq files were quality and adapter trimmed using trimmomatic in paired-end mode (ILLUMINACLIP:TruSeq3-PE.fa:2:30:10:8:true, AVGQUAL:25, MINLEN:20) (87). Unique molecular identifiers (UMIs) were extracted from paired-end reads by UMI-tools (v. 1.0.1) using the umi_tools command (--bcpattern=NNNN, --bc-pattern2=NNNN). Mapping was performed by bowtie2 (v. 2.2.5) in --sensitive-local mode with the last four nucleotides of each read being trimmed using --trim3 4 (88). SAM files were converted to BAMs, sorted and indexed using samtools (v. 1.9) (84). Subsequently, mapped reads were deduplicated using the extracted UMIs by umi_tools dedup with default settings for paired-end data. Detection of enriched 3’ ends by peak calling and downstream analysis was performed as described in (89). Briefly, strand-specific bedgraph files were first generated and CPM normalised using deepTools bamCoverage (v. 3.5.0), with the SAM flags 83 and 99 in combination with --Offset 1 and --binSize 1 -- minMappingQuality 20. Next, the termseq_peaks script was used with default parameters to call peaks using four replicates as input (https://github.com/nichd-bspc/termseq-peaks) (90). For end detection, peak files were merged with the position-specific count files using bedtools intersect (option -wao). Enriched positions were finally filtered and annotated based on the following criteria: For each peak the position with the highest number of reads was selected per replicate and only maximum peak positions selected that were present in at least three of the four replicates. Positions with less than five CPM counts were excluded from the analysis. Positions were assigned based on their relative orientation to a gene and their respective peak height as primary (highest peak within 300 bases downstream from a gene), secondary (each additional peak 300 bases downstream from a gene) and internal (each peak in the coding range).

### Nanopore sequencing

*Library preparation and sequencing:* For PCR-cDNA sequencing, ribosomal RNAs were first depleted using the Pan-Archaea riboPOOL (siTOOLs) according to the manucfactureŕs instructions with 1 µg DNase-treated input RNA (control condition, 95°C, mid-exponential growth phase). To prevent secondary structure formation and inefficient RNA treatment, samples were heat-incubated at 70°C for 3 minutes and immediately placed on ice before ligation. Next, a custom 3’ adapter (5ʹ-rAppCTGTAGGCACCATCAAT–NH2-3ʹ, NEB) was ligated to all RNAs following the protocol described in (91). This protocol is an alternative to the otherwise necessary enzymatic polyadenylation and improves 3’ end accuracy (92). Briefly, 100 ng rRNA-depleted RNA was mixed with 50 pmol 3’ adapter, 2 μl 10× T4

RNA ligase reaction buffer (NEB), 10 μl 50% PEG 8000 (NEB), 1 μl 40 U/μl−1 RNase Inhibitor (NEB), and 1 μl T4 RNA ligase 2 (truncated K227Q, NEB, 200,000 units/ml) and incubated at 16°C for 14 h. Finally, RNAs were cleaned up using the ZYMO RNA Clean and Concentrator kit (ZYMO). PCR-cDNA sequencing libraries were prepared following the Oxford Nanopore Technologies (ONT) PCR-cDNA barcoding kit protocol (SQK- PCB109) with minor modifications, such as using a custom 3’ cDNA RT primer (5ʹ- ACTTGCCTGTCGCTCTATCTTCATTGATGGTGCCTACAG-3ʹ, 2 µM), which replaces the VN primer. RNA and cDNA sizes were assessed using a Bioanalyzer (Agilent). After reverse transcription and template-switching as described in the ONT protocol, samples were PCR-amplified for 12 cycles with a 500 s elongation time. During library preparation, samples were quantified and quality checked using standard spectroscopic measurements (Nanodrop One, Thermo Fisher Scientific) and Qubit assays (Thermo Fisher Scientific). Equimolar amounts of samples were pooled, adapter-ligated, and sequenced on an R9.4 flow cell (Oxford Nanopore Technologies) using an MK1C device for 72 hours.

PCR-cDNA sequencing was performed for two biological replicates of cells grown to mid-exponential growth phase at 95°C.

### Data analysis of PCR-cDNA libraries

Basecalling of fast5 files was performed using guppy (v. 6.4.2+97a7f06) in high-accuracy mode (dna_r9.4.1_450bpd_hac.cfg) with standard parameters and a quality score threshold of 9. The PCR-cDNA library was demultiplexed by guppy_barcoder using default settings (–barcode_kits SQK-PCB109), except for disabling barcode trimming. Full-length sequenced reads were identified, strand-oriented, and trimmed using pychopper (v. 2.7.2, https://github.com/epi2me-labs/pychopper) with autotuned cutoffs and the recommended edlib backend for identifying custom primers. Cutadapt (v. 4.2) was used to remove remaining 3’-adapter sequences with the parameters -a “CTGTAGGCACCATCAAT” -j 0. Next, trimmed reads were mapped using minimap2 (v. 2.24-r1122) with standard parameters suggested for aligning Nanopore genomic reads (-ax map-ont) to the *P. furiosus* DSM 3638 genome (NCBI: CP023154.1) (93, 94). Alignments with more than 5 clipped bases (soft or hard clips) were removed using samclip (v. 0.4.0), and SAM files were converted to sorted BAM files using samtools (v. 1.16.1) (84).

Lastly, strand-specific coverage files were created using samtools depth (-a, -J options enabled) with a binsize of 1 and including reads with deletions in the coverage computation. Downstream analysis, including CPM normalization, calculating the mean coverage for each position of the two replicates, and plotting, was performed using the Tidyverse in R (95).

### Mass spectrometry

*Sample preparation, analysis and data processing:* Mass spectrometry analysis was performed for four biological replicates according to the following protocol. Protein samples were first purified by running them a short distance into a 4-12% NuPAGE Novex Bis-Tris Minigel (Invitrogen), followed by Coomassie staining and in-gel digestion with trypsin (96).

For generation of a peptide library, equal amount aliquots from each sample were pooled to a total amount of 200 µg, and separated into twelve fractions using a basic pH reversed phase C18 separation on an FPLC system (Äkta pure, Cytiva) and a staggered pooling scheme. All samples were spiked with a synthetic peptide standard used for retention time alignment (iRT Standard, Schlieren, Schweiz).

Protein digests were analyzed on a nanoflow chromatography system (nanoElute) hyphenated to a hybrid timed ion mobility-quadrupole-time of flight mass spectrometer (timsTOF Pro, all Bruker). In brief, 400 ng equivalents of peptides were dissolved in loading buffer (2% acetonitrile, 0.1% trifluoroacetic acid in water), enriched on a reversed-phase C18 trapping column (0.3 cm × 300 µm, Thermo Fisher Scientific) and separated on a reversed-phase C18 column with an integrated CaptiveSpray Emitter (Aurora 25 cm × 75 µm, IonOpticks) using a 60 min linear gradient of 5-35 % acetonitrile/0.1% formic acid (v:v) at 300 nl min^-1^, and a column temperature of 50⁰C.

DDA analysis was performed in PASEF mode (97) with 10 PASEF scans per topN acquisition cycle. Multiply charged precursors were selected based on their position in the *m/z*–ion mobility plane and isolated at a resolution of 2 Th for *m/z*≤700 and to 3 Th for *m/z*>700 for MS/MS to a target value of 20,000 arbitrary units. Dynamic exclusion was set to 4 min. Two technical replicates per C18 fraction were acquired.

DIA analysis was performed in diaPASEF mode (98) using 32×25 Th isolation windows from *m/z* 400 to 1,200 to include the 2+/3+/4+ population in the *m/z*–ion mobility plane. The collision energy was ramped linearly as a function of the mobility from 59 eV at 1/K0=1.6Vs cm^−2^ to 20 eV at 1/K0=0.6Vs cm^−2^. Two technical replicates per biological replicate were acquired.

Protein identification and quantification were performed in Spectronaut Software 15.6 (Biognosys). Protein identification was achieved using the software’s Pulsar algorithm at default settings against the UniProtKB *Pyrococcus furiosus* reference proteome (revision 09-2021) augmented with a set of 51 known common laboratory contaminants. For quantitation, up to the 6 most abundant fragment ion traces per peptide, and up to the 10 most abundant peptides per protein were integrated and summed up to provide protein area values. Mass and retention time calibration as well as the corresponding extraction tolerances were dynamically determined. Both identification and quantification results were trimmed to a False Discovery Rate of 1% using a forward-and-reverse decoy database strategy. Protein quantity distributions were normalized by quartile normalization, and intensity-based absolute quatification (iBAQ) values as proxies for protein levels (99).

#### Differential enrichment analysis

Quantitative values were processed using log_2_ transformation, and PCA was used for quality control and detection of outlier replicates (CS 2 replicate 1, CS 3 replicate 4, CS R replicate 2, HS 2 replicate 3, HS R replicate 1) after visual inspection. Normalized protein expression data was analyzed using the R limma package to identify differentially expressed proteins between control and cold or heat-stressed samples (100).

### Additional bioinformatic analysis

#### Normalisation

Total transcripts per million (TPM) was used as a normalization method for RNA-seq data analysis to account for differences in sequencing depth and transcript length. TPM values were calculated by dividing the number of reads mapping to each gene by the gene length in kilobases, then dividing the resulting reads per kilobase (RPK) values by the sum of all RPK values in the sample, and finally multiplying the quotient by one million. This normalization method ensures that the sum of all TPM values in each sample is the same, allowing for better comparison of gene expression levels between samples.

#### arCOG enrichment

Functional enrichment analysis based on the Archaeal Clusters of Orthologous Genes (arCOG) classification was performed as described previously (101, 102). Briefly, arCOGs for *P. furiosus* were retrieved from (51) and gene set enrichment analysis performed with the goseq package in R, which accounts for gene lengths bias (103). For each comparison, a condition- and method-specific background file was generated from all genes that could be detected. Next, p-values for overrepresentation of arCOG terms in the differentially expressed genes were calculated separately for up- and downregulated genes based on RNA-seq and MS data, respectively, and were considered as significantly enriched below a cutoff of 0.05.

#### Promoter and terminator analysis

Primary transcription start sites (TSS) and corresponding 5’ UTR lenghts were extracted from (47). Genes were categorized by the existence of an archaeal-typical promoter motif containing a TFB-recognition element (BRE, 5’- G/C(A/T)AAA-3’) and a TATA element (5’-TTT(A/T)(A/T)(A/T)-3’) (47, 104). Therefore, the sequences from −50 to +10 from the TSS were analysed via MEME (v. 5.4.1., -mod zoops -minw 8 -maxw 20) and the genes contributing to the motif categorized as + promoter (105). Position-specific motifs were plotted in R using the ggseqlogo package (106). Promoter strength was estimated according to the method used in (54). Briefly, sequences from −42 to −19 were extracted and analysed using MEME in oops mode. The p value from the motif search was used as an estimator for the promoter strength.

3’ ends were classified similarly based on the presence of a poly(U)- terminator motif. To this end, sequences from −35 to +2 from the primary 3’ ends were extracted and analysed via MEME (-mod zoops-minw 4 -maxw 20) and genes contributing to the motif classified as +poly(U) motif. Nucleotide enrichment analysis was performed as described in (55). The frequency of each base was calculated using the extracted sequences and compared to the same calculation based on randomly sampling 100,000 positions from intergenic regions. From this comparison the log2 was calculated and only the enrichment position of nucleotide U plotted. Structural stability of the RNA was predicted by folding the 45 nt long RNA upstream of the terminators using the LncFinder package in R, which uses the RNAfold software with standard parameters, and compared to 1,000 randomly subsampled intergenic regions (107).

#### Operon analysis

Operon analysis of selected heat-shock genes was performed by comparing the annotation from the DOOR2 operon database, the operon annotation based on ANNOgesic prediction using short-reads from mixed conditions from *P. furiosus* and from single-read analysis of PCR-cDNA Nanopore reads generated for this study (47, 108).

#### Correlation analysis

The Pearson correlation coefficient was used to calculate the correlation between values found in both samples (pairwise complete).

#### Detection and analysis of signature clusters

The z-score normalized log2 fold changes of RNA and protein values were subjected to principal component analysis (PCA) using the prcomp function in R. Next, the PCA results were used to calculate the pairwise Euclidean distances between samples. The resulting distance matrix was then used to perform hierarchical clustering with the ward.D2 linkage method. The number of clusters was determined using the elbow method and after inspection of enrichment analysis of arCOG categories. This approach allowed for the identification of clusters of genes with similar expression patterns across different conditions and datasets, which could provide insights into the regulation of cellular processes in the hyperthermophilic archaeon *Pyrococcus furiosus*.

Functional analysis was performed using the STRING database to identify known and predicted protein-protein interactions (PPIs) among the differentially expressed genes (109).

The codon adaption index (CAI) was computed using the CAI function from the coRdon package using the 5% most abundant proteins according to our control sample (110).

## Data availability

RNA sequencing data are available at the European Nucleotide Archive (ENA, https://www.ebi.ac.uk/ena) under project accession numbers PRJEB61174 (RNA-seq data used for differential gene expression analysis) and PRJEB61177 (Term-seq and Nanopore data) (111).

The mass spectrometry proteomics data have been deposited to the ProteomeXchange Consortium via the PRIDE partner repository with the dataset identifier PXD041262 (112).

## Code availability

Documentation and code of all essential analysis steps (tools and custom Rscripts) are available from https://github.com/felixgrunberger/HSCS_Pfu.

## Acknowledgements

We thank all the members of the Grohmann lab for fruitful discussions. H.U. and C.L. would like to acknowledge Lisa Neuenroth at the Department of Clinical Chemistry, University Medical Center Göttingen, for expert technical support of the mass spectrometric analyses. We furthermore would like to thank Eveline Peeters and Rani Baes for stimulating discussions and their expertise in archaeal biology.

## Supplementary Figures

**Fig. S 1.**
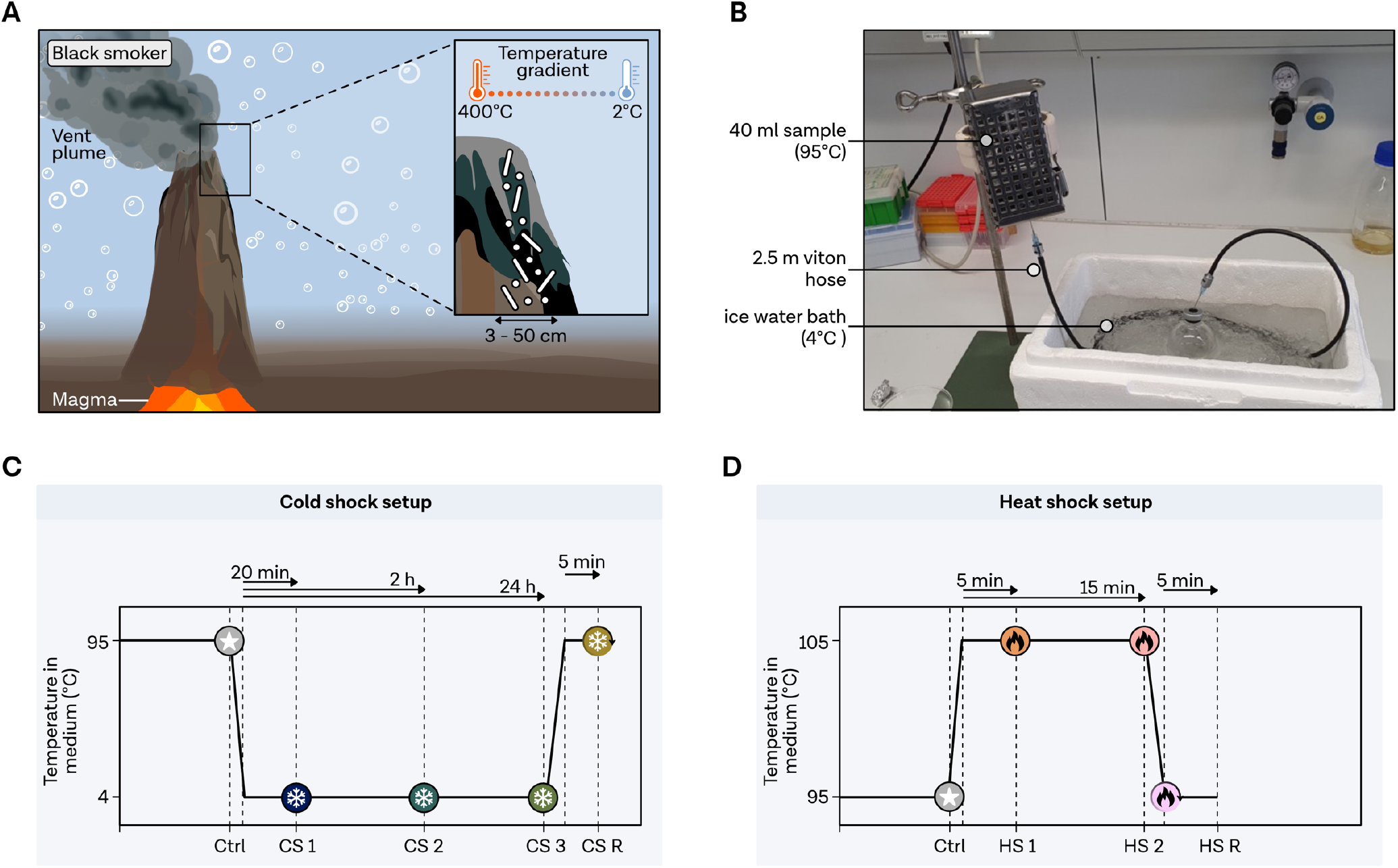
Natural and experimental temperature gradients for *Pyrococcus furiosus.* **A**, Black smokers, formed by effluents contacting magmatic heated rocks, are colonized by hyperthermophiles and characterized by large temperature gradients from 2°C to 400°C. Figure adapted from (12). **B**, Cold shock treatment setup: Mid-exponential phase cells underwent cold shock by rapidly cooling the medium through a 2.5 m long, 2 mm diameter viton hose flushed with anaerobic NaCl solution into a new bottle containing 1 bar excess nitrogen. **C**, Time points analyzed during cold and **D**, heat shock.

**Fig. S 2.**
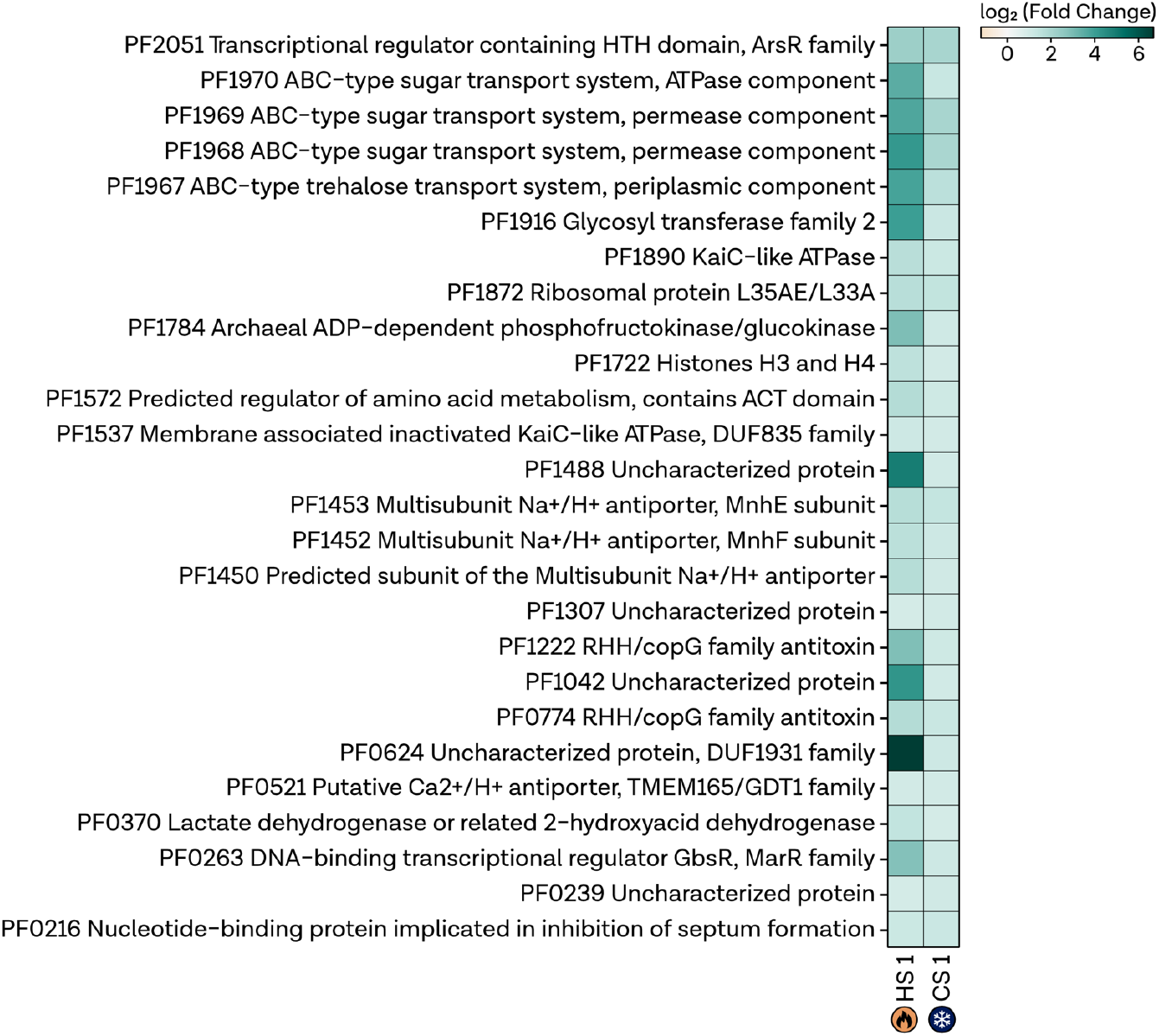
Fold-changes of genes upregulated during HS and CS. Comparison of RNA-seq fold changes between genes that are highly upregulated (>2-fold) during heat shock 1 (HS 1) and cold shock 1 (CS 1). Log_2_ fold change is indicated by the color bar.

**Fig. S 3.**
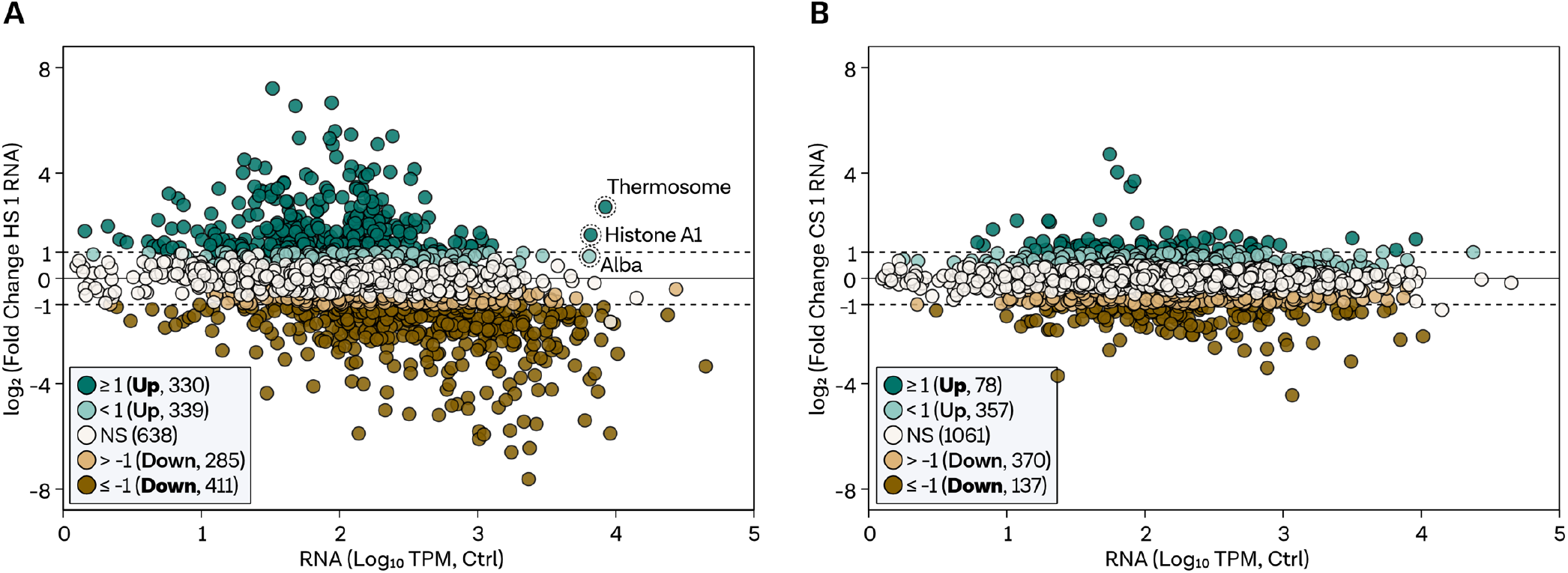
Comparison of expression values under control conditions to fold changes under stress conditions. **A**, Plot displaying log_2_ fold changes (y-axis) and TPM-normalized RNA counts (x-axis) comparing transcriptome changes at HS 1 (5 minutes) and **B**, CS 1 (20 minutes) to the expression levels at the control condition. Protein-coding transcripts are categorized by significance and fold changes, color-coded as strongly upregulated (padj < 0.05 & log_2_FC >= 1, dark green, bold font), upregulated (padj < 0.05 & log_2_FC < 1 & log_2_FC > 0, light green, normal font), non-regulated genes (NS, padj >= 0.05, white), strongly downregulated (padj < 0.05 & log_2_FC <= −1, dark brown, bold font), downregulated (padj < 0.05 & log_2_FC > −1 & log_2_FC < 0, light brown, normal font). Abundant HS genes are highlighted.

**Fig. S 4.**
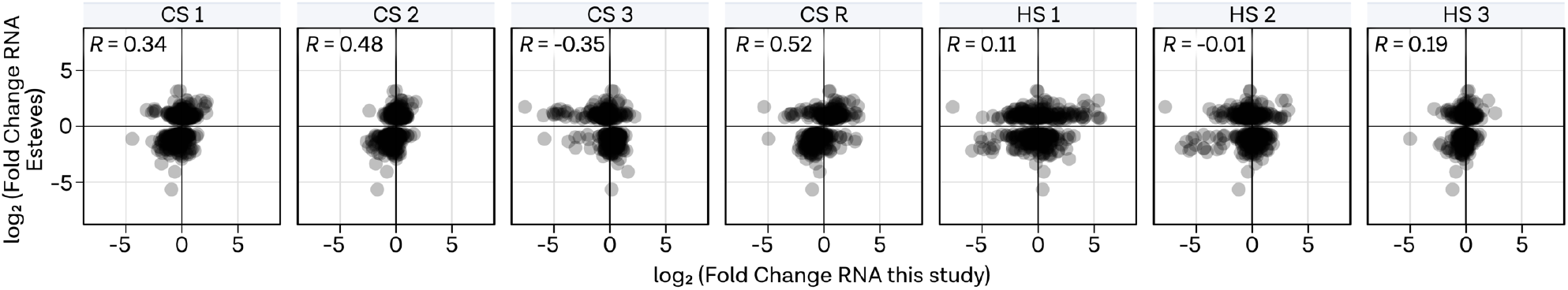
Comparison to fold changes from(113). Comparison of log_2_ fold changes of all stress conditions to the log_2_ fold changes observed in (113). Pearsońs correlation coefficient is shown in the top left for each comparison.

**Fig. S 5.**
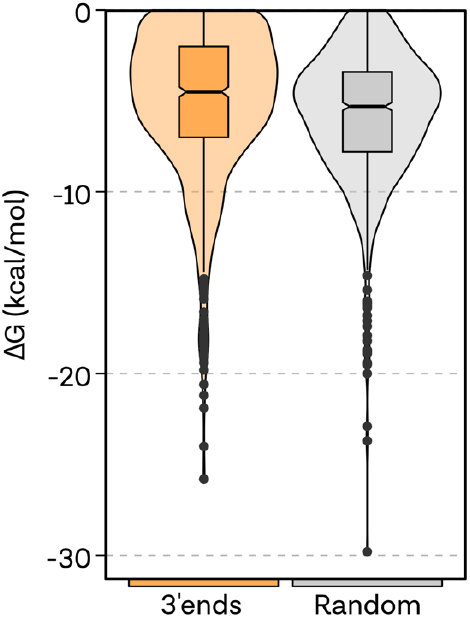
Secondary structure analysis. Distributions of predicted RNA structure stabilities for deteced 3’ ends (yellow) and random intergenic positions (grey).

**Fig. S 6.**
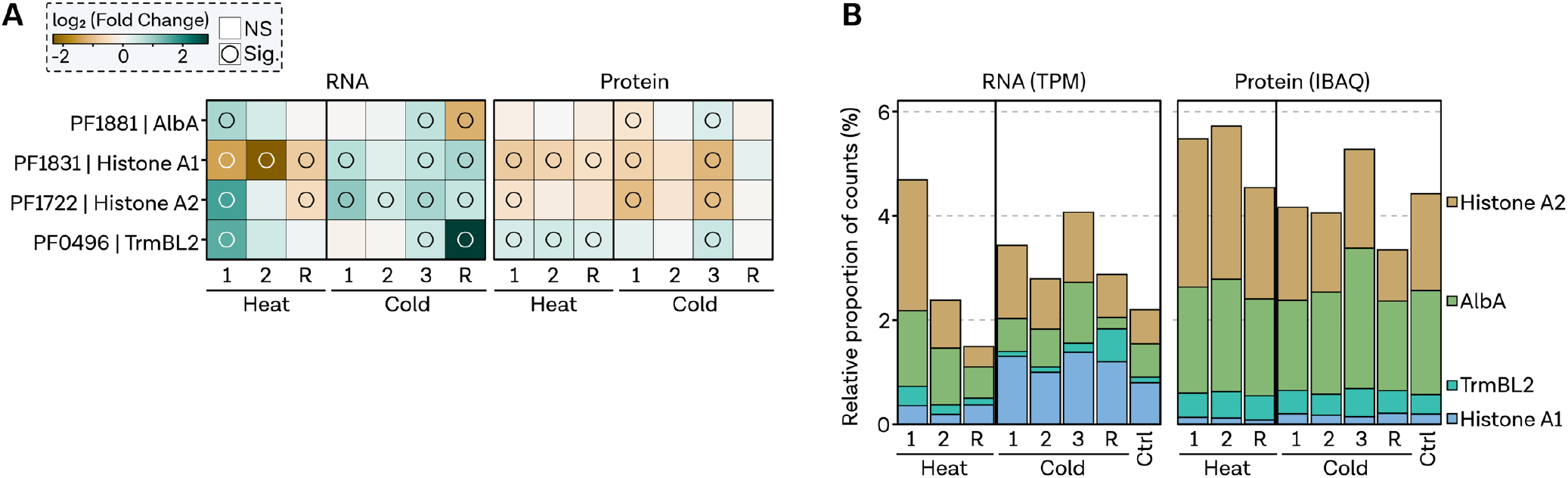
Overview of nucleoid-associated proteins (NAPs). **A,** Heatmap displaying log_2_ fold changes of NAPs, color-coded by fold change. Presence of a circle indicates gene significance with an adjusted p-value < 0.05. **B**, Relative proportion of counts calculated based on TPM and IBAQ values for all protein-coding genes.

**Fig. S 7.**
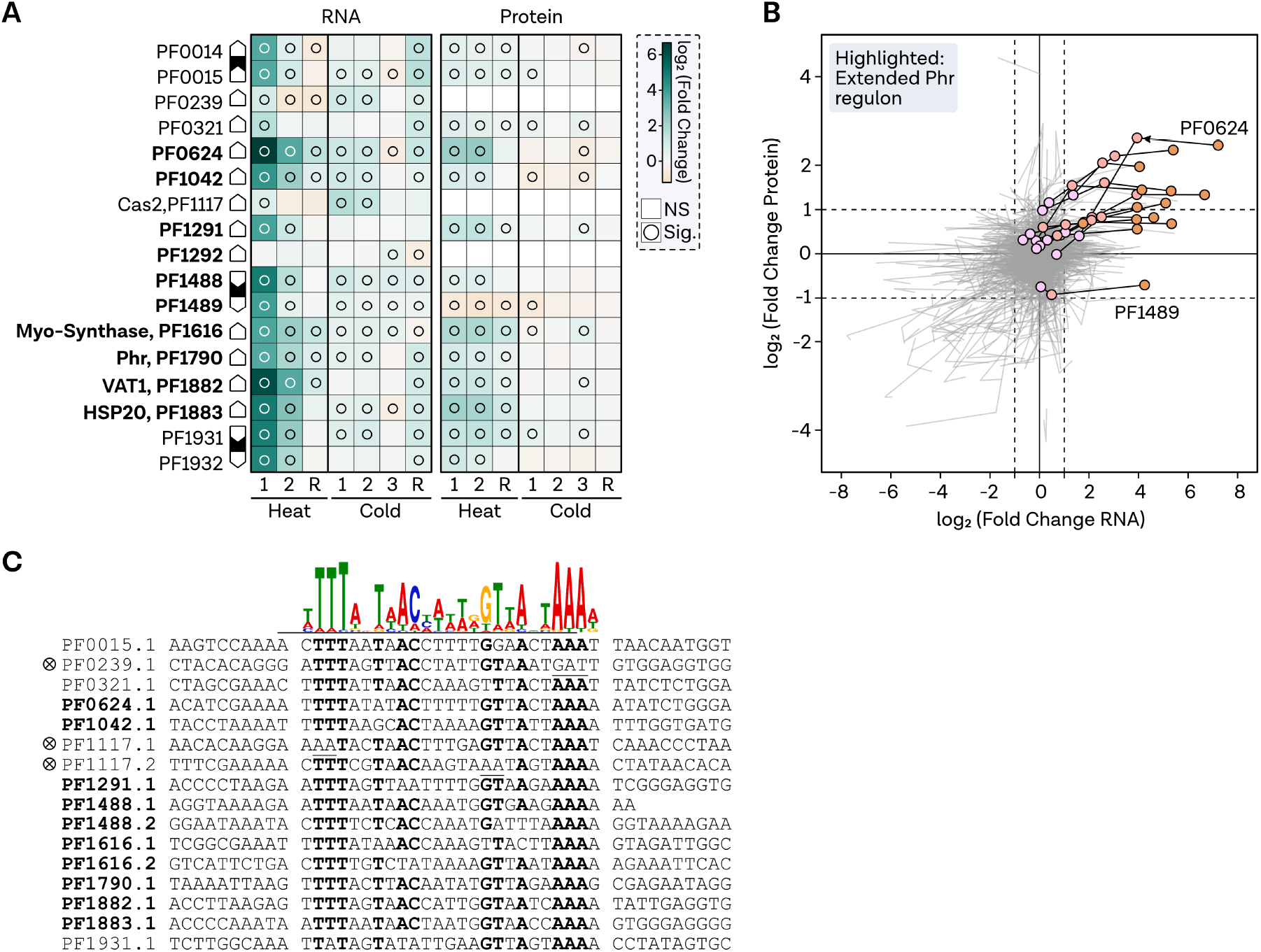
Analysis of extended Phr regulon. **A,** Heatmap displaying the log_2_ fold changes of all Phr targets, including experimentally verified or RegPrecise database-listed targets (40, 57). Fold change is color-coded, with circles denoting gene significance (adjusted p-value < 0.05). Operon organization is represented by connected rectangles. **B**, Pathway plot highlighting log_2_ fold changes in RNA (x-axis) and protein (y-axis) levels from HS 1 condition to HS 2 and recovery conditions. Extended Phr regulon genes are emphasized with colors and points. **C**, Multiple sequence alignment of upstream regions of Phr targets, displaying the consensus motif recognized by Phr at the top.

**Fig. S 8.**
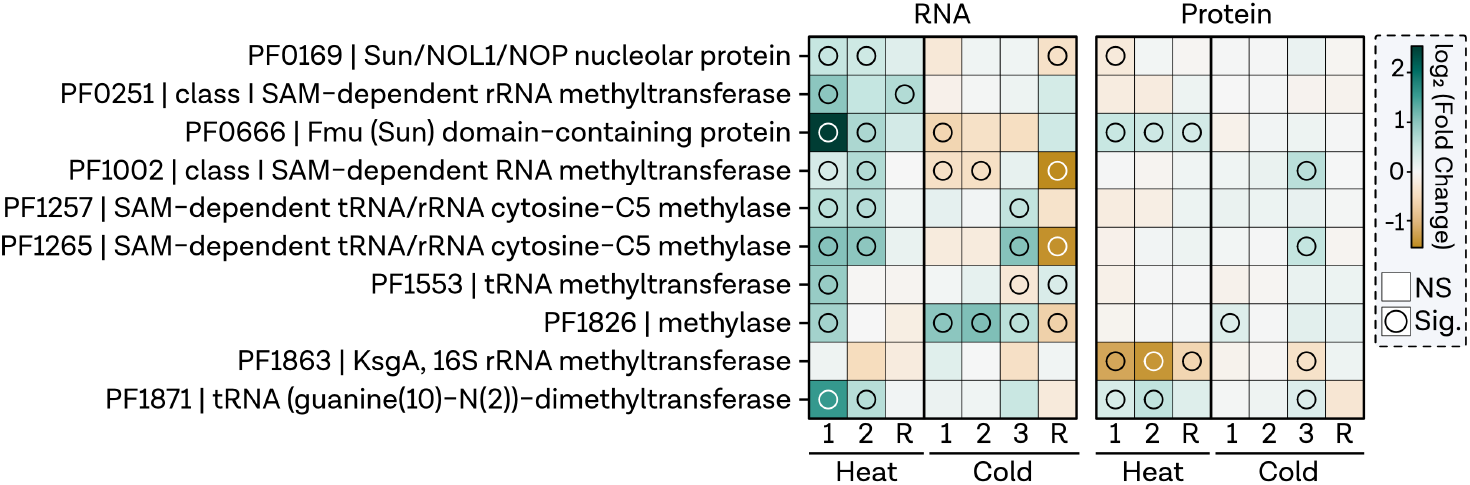
tRNA/rRNA-modifying enzyme gene expression. Heatmap illustrating log_2_ fold changes of genes annotated as tRNA/rRNA-modifying enzymes, color-coded based on log fold change. Circles represent gene significance (adjusted p-value < 0.05).

**Fig. S 9.**
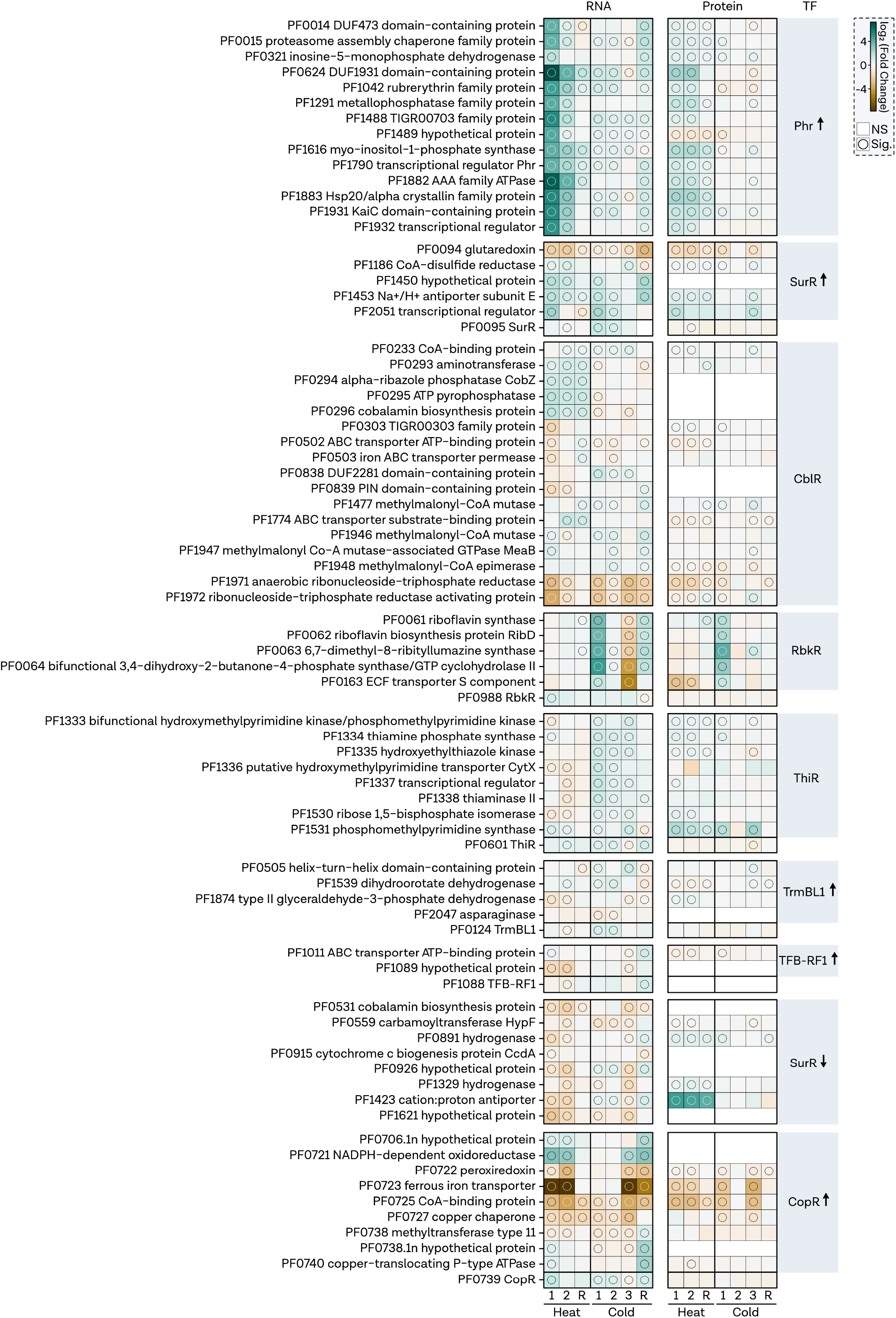
Overview of transcription factor regulon gene expression under heat and cold shock conditions. Heatmap showing log_2_ fold changes of genes within experimentally validated or predicted TF regulons, color-coded according to log fold change. Mode of TF is indicated by arrow up (activator) or arrow down (repressor). Circles indicate gene significance (adjusted p-value < 0.05).

**Fig. S 10.**
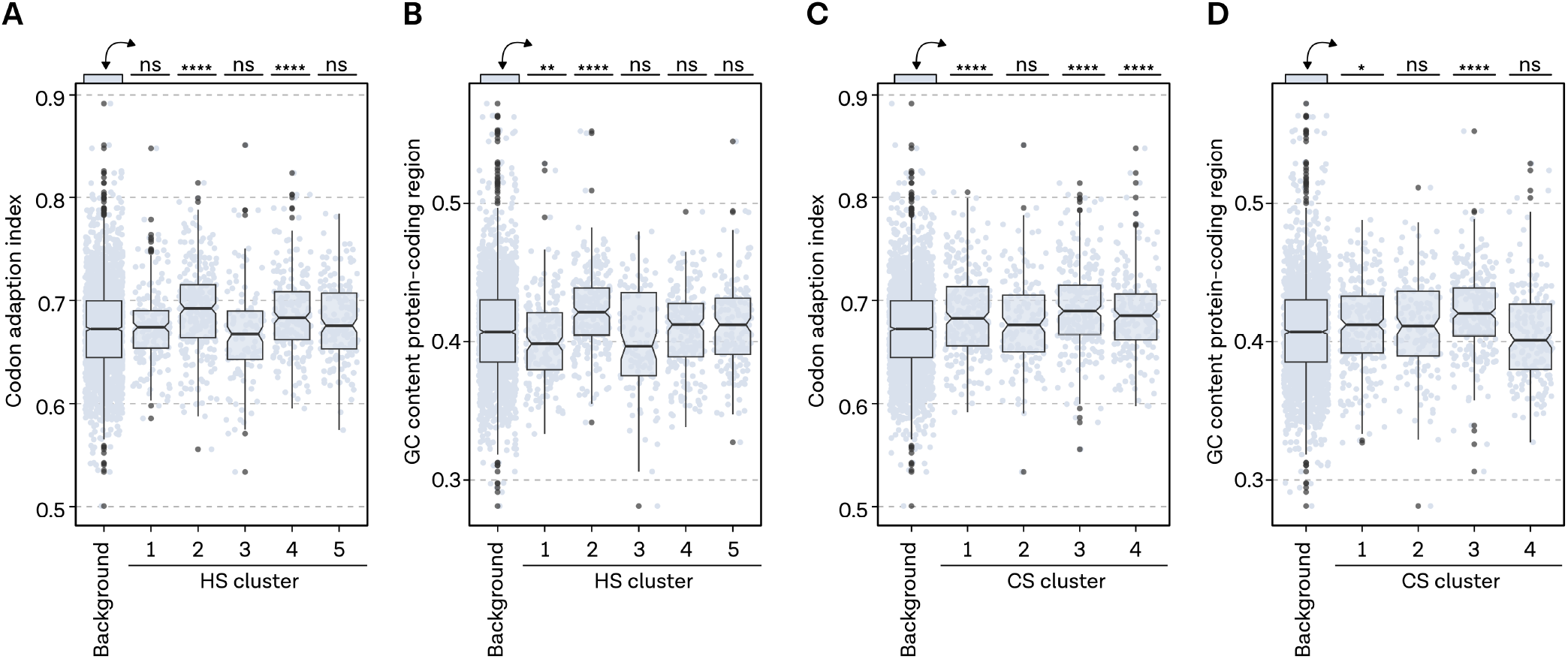
HS and CS cluster sequence features. **A,** Codon adaptation index distribution for background and HS clusters. Boxplot with edges indicating 1st and 3rd quartiles, median as center line, and whiskers denoting points within 1.5x interquartile range. Welch’s t-test assessed group differences; **** denotes p-values < 0.0001. **B**, GC content distribution in protein-coding regions. Boxplot with edges indicating 1st and 3rd quartiles, median as center line, and whiskers denoting points within 1.5x interquartile range. Welch’s t-test assessed group differences; ** denotes p-values < 0.01. **C**, Codon adaptation index distribution for background and CS clusters. Boxplot with edges indicating 1st and 3rd quartiles, median as center line, and whiskers denoting points within 1.5x interquartile range. Welch’s t-test assessed group differences; **** denotes p-values < 0.0001. **D**, GC content distribution in protein-coding regions for CS clusters. Boxplot with edges indicating 1st and 3rd quartiles, median as center line, and whiskers denoting points within 1.5x interquartile range. Welch’s t-test assessed group differences; * denotes p-values < 0.05.

